# CpG island density predicts CBP/p300 dependency across 3D chromatin clusters

**DOI:** 10.64898/2026.05.04.722036

**Authors:** Md Imdadul H. Khan, Matthew S. Chang, Yaw Asante, Maya K. Al-Haddad, Teagan Kukhta, Bhavatharini Udhayakumar, Ana María Garzón-Porras, Elena-Loredana Rotariu, Saanvi Aneja, Jordyn L. Kelly, Carrietta B. Farma-Hai, Kavi Ullal, Diana H. Chin, Alexander J. Federation, Lindsay K. Pino, Julia Robbins, Marco Wachtel, Shaun R. Stauffer, Berkley E. Gryder

## Abstract

RNA Polymerase II (Pol2) organizes transcription through higher-ordered chromatin clusters that integrate promoter and enhancer interactions to coordinate gene expression. Nevertheless, the features that distinguish unique classes of Pol2-mediated clusters remain to be defined. Here, we identify two distinct classes of Pol2-mediated clusters: one enriched for CpG islands, promoter-promoter interactions, and housekeeping gene expression, and another characterized by high CBP/p300 occupancy, enhancer-promoter looping, and lineage-defining (LD) transcriptional programs. Acute inhibition of CBP/p300 catalytic activity leads to rapid loss of acetylation at enhancers and preferential downregulation of LD genes, resulting in impaired cellular proliferation and activation of apoptotic programs. Integrative machine learning modeling reveals that cluster strength, RNA half-life, and CpG island content as strong predictors of genes sensitive to CBP/p300 inhibition. Together, these findings clarify enhancer-addiction and vulnerability to CBP/p300 inhibition.

## Introduction

Gene regulation is fundamentally governed by the selective activation of distal *cis*-regulatory elements known as enhancers. For distal communication to be maintained across the genome, chromatin must be organized into accessible and functionally distinct domains. Increasing evidence supports an organized model of chromatin in which regulatory elements and transcriptional machinery are spatially concentrated^1–3^.

At the center of this is RNA Polymerase II (Pol2), the twelve-subunit enzyme that facilitates transcription of all protein-coding genes^4^. In recent years, Pol2 has been observed to accumulate within discrete chromatin regulatory hubs, often referred to as clusters or transcriptional condensates, that connect multiple regulatory elements through dynamic looping interactions^5^. These clusters are shaped by a combination of histone post-translational modifications, enhancer and promoter density, and transcriptional co-activators, including the paralogous histone acetyltransferases (HATs) CREB-binding protein (CBP) and E1A binding protein p300 (p300). CBP/p300 play a central role by depositing acetyl groups to the tails of histones to promote chromatin accessibility and enhancer activity. However, recent studies demonstrate that acute catalytic inhibition of CBP/p300 can alter transcription without inducing changes in chromatin accessibility^6^. This suggests that some promoters can remain accessible and active independently of enhancer activity. One such feature that may help explain this is CpG islands: GC-rich DNA sequences commonly localized at the transcription start sites of promoters that are intrinsically accessible. The presence of these accessible promoters raises the possibility that Pol2-mediated clusters form around regulatory elements with distinct chromatin environments^7^. Despite the well-established role of CBP/p300 in transcription and cancer, it remains unclear whether Pol2-mediated clusters are uniform in their composition or are organized into distinct classes with differential dependence on CBP/p300 catalytic activity.

In this study, we used Pol2 HiChIP to define two distinct classes of Pol2-mediated clusters: one enriched for CpG islands, promoter-promoter interactions, and housekeeping (HK) gene expression, and another characterized by high CBP/p300 occupancy, enhancer-promoter looping, and expression of lineage-defining (LD) genes. To interrogate these dependencies mechanistically, we developed IHK-44, a potent inhibitor of the CBP/p300 HAT domain. Using IHK-44, we demonstrate that enhancer-associated acetylation, particularly histone H2B N-terminal lysine acetylation, undergoes rapid turnover and is selectively vulnerable to CBP/p300 inhibition. This vulnerability translates into preferential suppression of lineage-defining transcriptional programs, collapse of downstream protein expression, and selective growth arrest in enhancer-addicted cancers. By extending our analysis across PRISM’s 958 cancer cell lines^8^, we identify histone H2B hyperacetylation as a chromatin signature of enhancer dependency and a predictive marker of sensitivity to CBP/p300 HAT inhibition. Finally, machine learning models determined that cluster strength, RNA half-life, and CpG island content are strong predictors of gene response to CBP/p300 inhibition.

Together, our findings establish a unifying framework in which gene function, 3D chromatin architecture, and epigenetic dependency are mechanistically linked, revealing enhancer-dense regulatory domains as a selective therapeutic vulnerability in enhancer-addicted cancers.

## Results

### Features distinguishing housekeeping and lineage-defining 3D architectures

The spatial organization of the genome plays a decisive role in establishing and maintaining cell identity^7,9^. By folding into higher-ordered structures, gene promoters and distal enhancers are brought into physical proximity, forming regulatory hubs (hereby referred to as clusters) that coordinate gene expression programs^10–12^. However, it remains unclear whether these organizational principles apply uniformly across genes with distinct regulatory functions. We hypothesized that lineage-defining (LD) genes are often embedded within enhancer-rich clusters, whereas housekeeping (HK) genes that sustain basic cellular functions reside in promoter-rich clusters.

To test this hypothesis, we used RNA polymerase II (RNA Pol2) HiChIP to define chromatin interactions of active transcription connecting gene promoters and distal enhancers (**Figure 1a**). We used PAX3-FOXO1 (P3F) fusion-positive rhabdomyosarcoma (FP-RMS) as our model system because its chromatin architecture has been extensively characterized^13–15^. After ranking clusters by interaction strength and assigning a target gene to each cluster based on maximum transcript abundance across connected loci, we examined the structural and regulatory elements at cluster anchors. In general, chromatin is opened by the activation of enzymes (histone writers) or by the presence of naturally histone-poor DNA sequences (CpG dinucleotide-rich regions, or CpG islands)^16^. We previously discovered that P3F recruits the histone writer p300 to its C-terminal activation domain and thereby shapes Pol2 cluster formation (**Supplemental Figures S1a and S1b**)^17^. To determine whether CpG island content affects the organization of RNA Pol2-mediated clusters, we quantified CpG island overlap at cluster anchors and ranked clusters by CpG island content (**Figure 1b**). Notably, a broad spectrum of CpG island content was observed among clusters, with 359 clusters exhibiting high CpG island content (≥75% of anchors overlapping CpG islands), 304 clusters exhibiting medium CpG island content (>33% and <75%) and 349 clusters exhibiting low CpG island content (≤33% overlap). The composition of cluster features revealed systematic differences across this CpG island spectrum (**Figure 1c**). Clusters with high CpG island content were enriched for promoter-promoter (P-P) loops and were preferentially associated with HK genes. In contrast, clusters with low CpG island content showed greater occupancy by p300 and P3F and were enriched for LD genes. Consistent with this, we quantified CpG island content by gene category and found that HK gene clusters exhibited the highest levels of CpG island content while LD gene clusters showed markedly lower CpG island content, suggesting that CpG content is a fundamental distinguisher of cluster function (**Figure 1d**).

**Figure 1.**
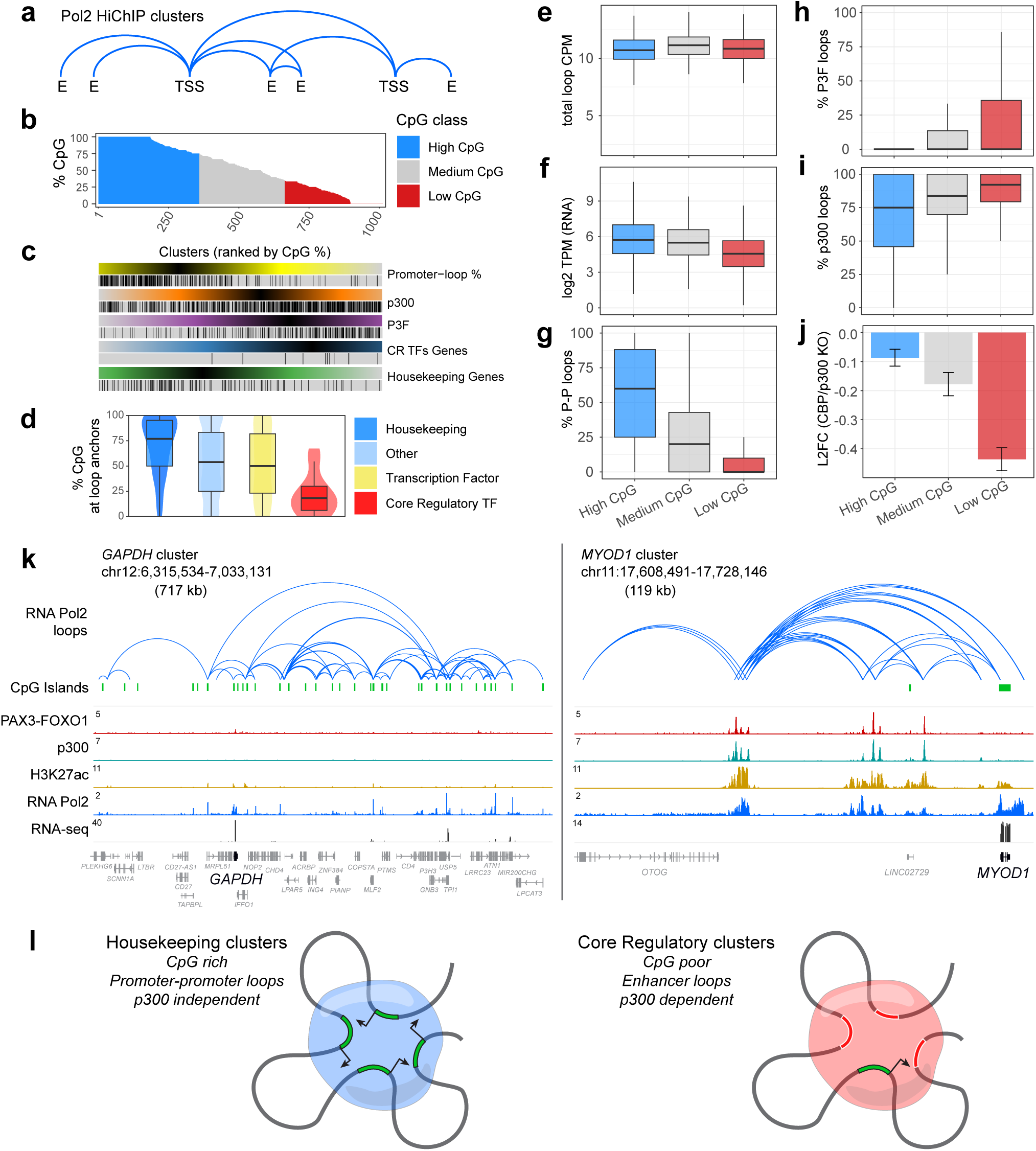
Features distinguishing housekeeping from lineage-defining clusters. **a.** Overview of Pol2 HiChIP derived clusters. **b.** Clusters ranked by the percentage of their loop anchors overlapping CpG islands, broken into high, medium and low CpG content. **c.** Overlap of cluster features, including promoter-promoter loops, p300 and PAX3-FOXO1 ChIP-seq peak overlap, and target gene categories (Core Regulatory Transcription Factors and Housekeeping gene), at clusters ranked by percent CpG islands at anchors. **d.** Percentage of CpG islands at loop anchors within clusters, separated by gene category. **e.** Total Pol2 contacts per million (CPM) signal among clusters grouped by CpG content. **f.** Gene expression from RNA-seq (log2 transcripts per million, TPM) among clusters grouped by CpG content. **g.** Percentage of promoter-promoter loops among clusters grouped by CpG content. **h.** Frequency of PAX3-FOXO1 binding at loop anchors in clusters grouped by CpG content. **i.** Frequency of p300 binding at loop anchors in clusters grouped by CpG content. **j.** Gene expression changes in RMS cells after dual CRISPR mediated knockout of CBP and EP300, for genes in clusters grouped by CpG content. **k.** GAPDH and MYOD1 gene regulatory clusters, shown with Pol2 HiChIP clustering, CpG island abundance, P3F and p300 binding, H3K27ac and Pol2 ChIP-seq levels, and RNA-seq, revealing distinct cluster features. **l.** Model of CpG rich Housekeeping clusters vs. p300 rich Core Regulatory clusters.

We next asked whether clusters with different CpG island densities varied in their transcriptional output and loop composition. Despite differences in CpG island and p300 distribution, overall RNA Pol2 density was comparable across cluster types (**Figure 1e**). Gene expression levels were moderately higher in CpG-rich clusters than in CpG-poor clusters (Fi**gure 1f**). As expected, CpG-rich clusters contained greater P-P loop interactions compared to CpG-poor clusters (**Figure 1g**). P3F binding events were rare in CpG-rich clusters but increased progressively with lower CpG content (**Figure 1h**). p300 occupancy was widespread across clusters but showed the greatest enrichment in CpG-poor clusters (**Figure 1i**). Genes residing in CpG-poor clusters exhibited the greatest reduction in gene expression levels following CBP/p300 loss, whereas genes in CpG-rich clusters showed minimal loss (**Figure 1j**).

Examples illustrating CpG-rich and CpG-poor clusters are shown for the HK gene glyceraldehyde-3-phosphate dehydrogenase (*GAPDH*) and the lineage-defining myogenic differentiation 1 gene (*MYOD1*), respectively (**Figure 1k**). Together, these data support a model in which CpG-rich clusters form P-P Pol2 loops that drive constitutive transcription largely independent of p300, while CpG-poor clusters rely on p300-dependent enhancer-mediated Pol2 loops characteristic of LD regulatory programs (**Figure 1l**).

### IHK-44 inhibits the HAT domain of CBP/p300 and reduces histone acetylation

CBP/p300 are homologous histone acetyltransferases (HATs) that open chromatin by depositing acetyl marks on lysine residues of histone tails. Their catalytic activity is essential for establishing environments of high enhancer activity and transcriptional output. Having discovered that enhancer-rich clusters are marked by high p300 content, we hypothesized that they must be particularly sensitive to pharmacological inhibition of CBP/p300. Yet, CBP/p300 inhibitors available during the course of our studies lacked desirable levels of potency and selectivity, motivating us to search for improved chemical scaffolds to develop enhanced inhibitors of the HAT domain.

In 2022, a team at GlaxoSmithKline identified proline-based p300 inhibitors from a DNA-encoded library. Subsequent lead optimization introduced a pyrazolo[4,3-*b*]pyridine moiety and a cyclobutyl ring, with additional fluorination of both the proline and cyclobutyl rings^18^. This effort yielded compound 28, a highly potent p300 inhibitor that nonetheless exhibited metabolic liabilities *in vivo*. Concurrently, Daiichi Sankyo discovered 1H-indazole-based p300 inhibitors through high-throughput screening and applied scaffold hopping from a 1,4-oxazepane core to generate a lead series^19^ culminating in DS-9300, a potent CBP/p300 inhibitor with improved *in vivo* metabolic stability. Notably, DS-9300 is structurally similar to compound 28, differing primarily by replacement of the cyclobutyl ring with a cyclohexyl moiety and substitution of the chloro group of the phenyl ring with a methoxy group^20^. Co-crystal structure (PDB ID: 8GZC) revealed that the N4 nitrogen of the pyrazolo[4,3-b]pyridine core forms a water-mediated hydrogen bond with the NH of Tyr1446, while the N1–H engages in a water-mediated hydrogen bond with the backbone carbonyl oxygen of Pro1452. Asp1399 and Ser1400 also form key interactions with the amide NH and carbonyl oxygen, respectively. Computational analysis further indicated that the terminal phenyl group is solvent-exposed and conformationally flexible, lacking hydrogen-bonding or π–π interactions. Guided by these insights, we functionalized the phenyl terminus with variety of functional groups to develop IHK-44, a potent CBP/p300 inhibitor (**Figures 2a and 2b**).

**Figure 2.**
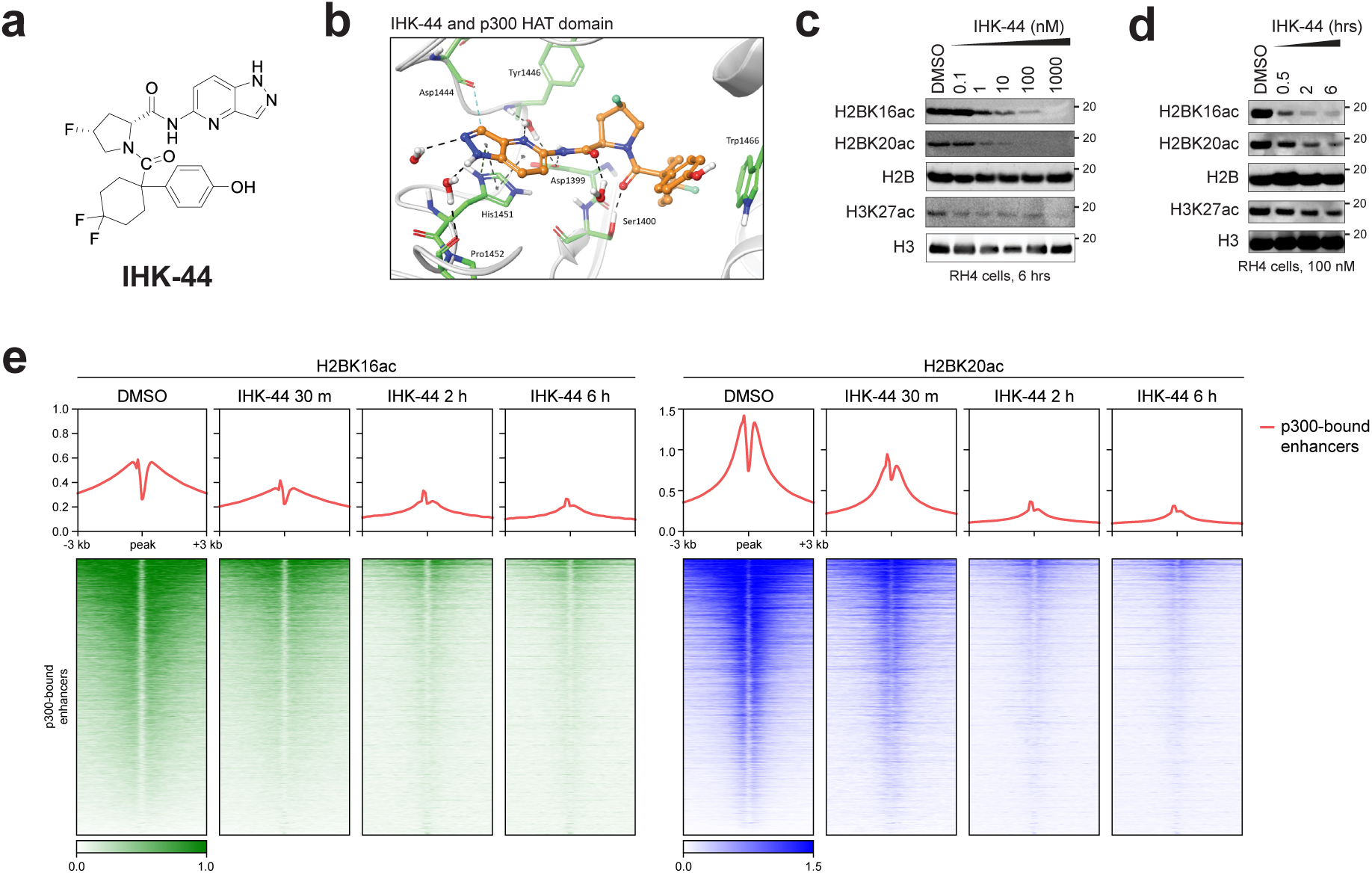
IHK-44 binds to the HAT domain of CBP/p300 and reduces histone acetylation. **a.** Molecular structure of IHK-44. **b.** IHK-44 docked to the HAT domain of p300. **c-d.** Dose-response (**c**) and time-course (**d**) Western blot of RH4 cells after IHK-44 treatment. Dose-response samples were treated for 6 h each. Time-course samples were treated at 100 nM each. H2B and H3 are shown as loading controls. **e.** Heatmaps of HBK16ac and H2BK20ac ChIP-seq shows that p300-bound enhancers are downregulated after 100 nM IHK-44 treatment over a 6 h time-course treatment period.

We first measured the ability of IHK-44 to reduce histone acetylation. Although H3K27ac is a canonical marker for active enhancers and promoters, H2B N-terminus lysine acetylation (H2BNTKac) is a more direct readout of CBP/p300 catalytic activity^21,22^. H2BNTKac marks, particularly H2BK16ac and H2BK20ac, are enriched at active enhancers, making them direct readouts of enhancer-rich clusters. Treatment with IHK-44 for 6 hours in a dose-dependent manner displayed a pronounced reduction in H2BK16ac and H2BK20ac compared to H3K27ac (**Figure 2c**). Time-course treatment revealed that these marks were rapidly reduced, whereas H3K27ac remained less changed even after 6 hours (**Figure 2d**).

Given this selective and rapid loss of H2BNTKac, we then asked how IHK-44 compares within the broader landscape molecules targeting CBP/p300. To test this, we benchmarked IHK-44 against compounds that interact with CBP/p300 trough different mechanisms. These included CBP/p300 bromodomain inhibitors CCS1477 (ref ^23^) and GNE-049 (ref^24^), the catalytic HAT inhibitor A-485 (ref^25^), the selective p300 degrader JQAD1 (ref^26^), and the CBP/p300 degraders dCBP-1 (ref^27^) and CBPD-409 (ref^28^). Western blot analysis of RH4 cells treated with each these compounds showed that IHK-44 reduced acetylation to levels comparable to CBP/p300 degraders (**Supplemental Figure 2**). The exception was JQAD1, whose selective degradation of p300 but not CBP limited its impact. Bromodomain inhibitors failed to suppress acetylation, consistent with reports that they only affect a subset of CBP/p300-dependent acetylation events^29^. A-485 achieved only modest suppression when compared to IHK-44. Together, these findings position IHK-44 as an effective inhibitor of CBP/p300 catalytic activity.

We next asked where this catalytic inhibition manifests across the genome. To address this, we first defined the H2BNTKac landscape in FP-RMS using H2BK16ac and H2BK20ac ChIP-seq in RH4 cells. Under DMSO conditions, both H2BK16ac and H2BK20ac displayed minimal overlap at promoter-proximal regions and CpG islands (**Supplemental Figures 3a and 3b**). In contrast, both markers were highly enriched at promoter-distal regions and p300 sites, consistent with the idea that CBP/p300 deposits H2BNTKac at enhancers (**Supplemental Figures 3a and 3c**). Notably, H2BK16ac and H2BK20ac overlapped ATAC-seq sites and greatly with H3K27ac sites, consistent with previous studies that H3K27ac acetylates active promoters and enhancers (**Supplemental Figures 3d and 3e**). This enhancer-rich distribution raised the possibility that H2BNTKac is not merely a general acetylation readout but rather encodes the enhancer hierarchy that defines super enhancers (SEs) in FP-RMS. Indeed, H2BK16ac-defined SEs displayed 4.6-fold greater enrichment than typical enhancers (TEs), including previously defined FP-RMS SEs like *MYOD1*, *JUN*, and *MEST* (**Supplemental Figures 3f and 3g**). H2BK20ac-defined SEs showed the same trend, with SEs showing 4.1-fold greater enrichment than TEs (**Supplemental Figures 3h and 3i**). Examples of gene tracks illustrating the H2BNTKac SE at *MYOD1* are shown (**Supplemental Figure 3j**).

**Figure 3.**
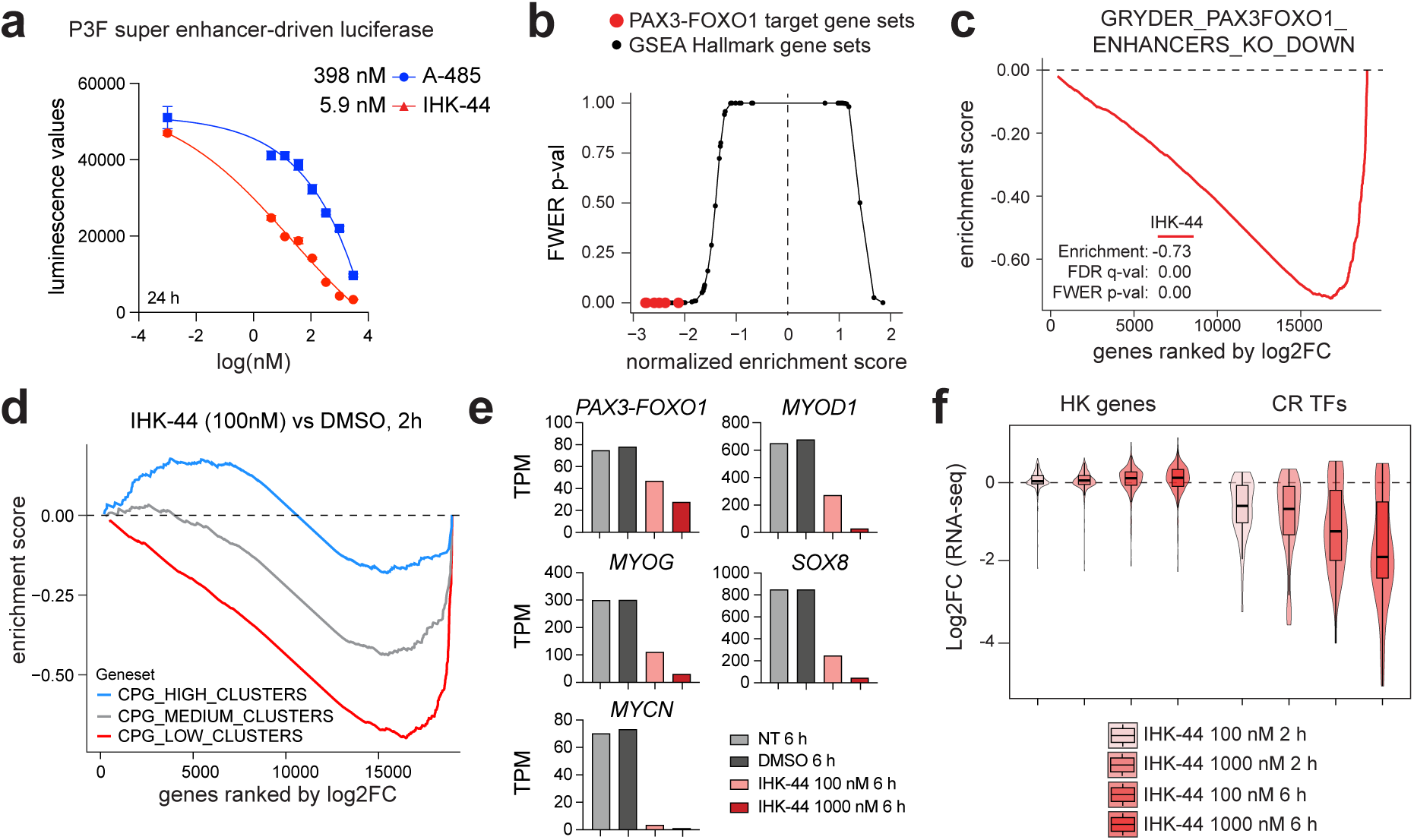
IHK-44 potently and selectively downregulates P3F target genes. **a.** Dose-response curves of super-enhancer-driven luciferase reporter activity following treatment with A-485 (blue) and IHK-44 (red) after 24 h. IHK-44 has a lower IC_50_ value (5.9 nM) than A-485 (398 nM). Values represent mean ± SEM of *n = 3* technical replicates. **b.** Scatter plot of FWER p-value versus normalized enrichment score (NES) of the MSigDB Hallmark gene sets and curated PAX3-FOXO1 target gene sets in RH4 cells treated with 100 nM IHK-44 for 6 h. PAX3-FOXO1 target gene sets are highlighted in red. **c.** GSEA analysis of genes downregulated after PAX3-FOXO1 knockout in RH4 cells treated with 100 nM IHK-44 for 6 h. **d.** GSEA analysis of genes downregulated with high (blue), medium (gray), and low (red) CpG island content after 100 nM IHK-44 treatment for 2 h in RH4 cells. **e.** TPM values *PAX3-FOXO1, MYOD1, MYCN, MYOG,* and *SOX8* after increasing IHK-44 treatment for 6 h in RH4 cells. **f.** Degree of downregulation of housekeeping (HK) genes and core regulatory transcription factors (CR TFs) after IHK-44 treatment in RH4 cells. Box plot represents median and quartiles, whiskers showing 1.5 × inter-quartile ranges.

We next asked whether IHK-44 selectively disrupts enhancer-associated acetylation. We performed ChIP-seq for H2BK16ac and H2BK20ac with 100 nM IHK-44 for 30 minutes, 2 hours, and 6 hours, and compared these profiles to DMSO-treated controls. Indeed, IHK-44 induced a rapid loss of H2BK16ac and H2BK20ac at p300-bound enhancer sites (**Figure 2e**). These results suggested that enhancer elements differed not only in their magnitude of acetylation loss but also in the rate in which acetylation is erased following CBP/p300 inhibition. We reasoned that the time-dependent loss of H2BNTKac could be approximated using a pseudo-first order kinetics framework. In this model, spike-in normalized H2BK16ac and H2BK20ac ChIP-seq signal was quantified across time for each DMSO peak and fitted to an exponential decay model to estimate a decay rate (*k_loss_*) and corresponding half-life (*t_1/2_*) (**Supplemental Figure 4a**). This revealed a broad distribution of acetylation half-life across all regulatory elements (**Supplemental Figure 4b**). When stratified by genomic feature, enhancers elements showed significantly shorter half-lives than promoter-proximal regions (**Supplemental Figure 4c**). Notably, promoters that overlapped CpG islands had greater half-lives than promoters that did not, suggesting that HK genes are more resistant to reduction of histone acetylation by CBP/p300 (**Supplemental Figure 4d**). Enhancers that co-occupied p300 showed shorter half-lives than enhancers that did not co-occupy p300, supporting the dependency of H2BNTKac with CBP/p300 occupancy (**Supplemental Figure 4e**). Regions that overlap LD genes showed lower half-lives than regions that overlap HK genes (**Supplemental Figure 4f**). Examples of gene tracks depicting this are shown for the LD gene, SRY-box transcription factor 8 (*SOX8*) (**Supplemental Figure 4g**). Together, these data support the use of IHK-44 as an effective chemical to probe enhancer function.

### IHK-44 selectively downregulates lineage specific genes and proteins

Given our observation that IHK-44 preferentially disrupts enhancer-mediated acetylation, we hypothesized that this would translate into a selective inhibition of enhancer-driven gene expression. To test this, we used a luciferase reporter that outputs P3F-driven transcription through a super enhancer regulatory element^30^. We observed that IHK-44 (IC_50_ = 5.9 nM) inhibits the reporter 67-fold more than A-485 (IC_50_ = 398 nM) (**Figure 3a**). To define what transcriptional programs were downregulated, we used RNA-seq and observed that P3F target gene sets were all statistically significant and negatively enriched, whereas Hallmark gene sets had a range of enrichments and statistical significance, suggesting selectivity was limited to P3F targets (**Figure 3b**). Genes previously defined as P3F- or CBP/p300-dependent were among the most strongly downregulated (**Figure 3c and Supplemental Figure 5a**). Importantly, genes in Pol2 clusters with low CpG island content were more sensitive to IHK-44 treatment than genes in clusters with medium or high CpG island content (**Figure 3d**). In particular, genes known for their critical roles in FP-RMS identity, including *P3F*, *MYOD1*, myogenin (*MYOG*), *SOX8*, and N-Myc proto-oncogene (*MYCN*), showed a dose-dependent reduction, with a near-complete loss of mRNA at 1 µM (**Figure 3e**). IHK-44 selectively downregulated LD genes in a time- and dose-dependent manner without affecting HK genes, consistent with the idea that enhancer-rich, CpG-poor clusters are selectively vulnerable to CBP/p300 inhibition while CpG-rich, promoter-centric clusters are resistant (**Figure 3f**).

To benchmark IHK-44 against other CBP/p300 probes, we performed RNA-seq on A-485, dCBP-1, and JQAD1. Principal component analysis of RNA-seq profiles revealed that IHK-44 clustered closely with dCBP-1 and diverged from A-485 over time (**Supplemental Figure S5b**). Interestingly, JQAD1 remained near NT and DMSO samples, suggesting that loss of p300 alone does not reduce target gene expression of FP-RMS cells and instead relies on CBP to compensate for p300 loss. Consistent with this, gene expression of FP-RMS CR TFs revealed a dose- and time-dependent loss by IHK-44 and dCBP-1, partial suppression with A-485, and minimal effects with JQAD1 (**Supplemental Figure S5c**).

Proteome level changes are closer to biology than RNA-seq, but many LD genes produce low abundance TFs that are difficult to measure with traditional proteomics. Therefore, we applied chromatin enriching salt separation coupled to data independent acquisition (CHESS-DIA) mass spectrometry to quantify protein-level response to IHK-44 (**Figure 4a**) ^31^. Fractionation of the nuclear compartments of RH4 cells confirmed that CBP and p300 are predominantly enriched in euchromatin fraction (**Figure 4b**). Proteomic analysis of euchromatin fractions following treatment with increasing doses of IHK-44 displayed that CBP/p300 protein abundance bound to chromatin remain largely unchanged (**Figure 4c**). However, the abundance of downstream lineage-defining proteins, such as P3F, MYOD1, and MYOG, were significantly reduced (**Figure 4d**).

**Figure 4.**
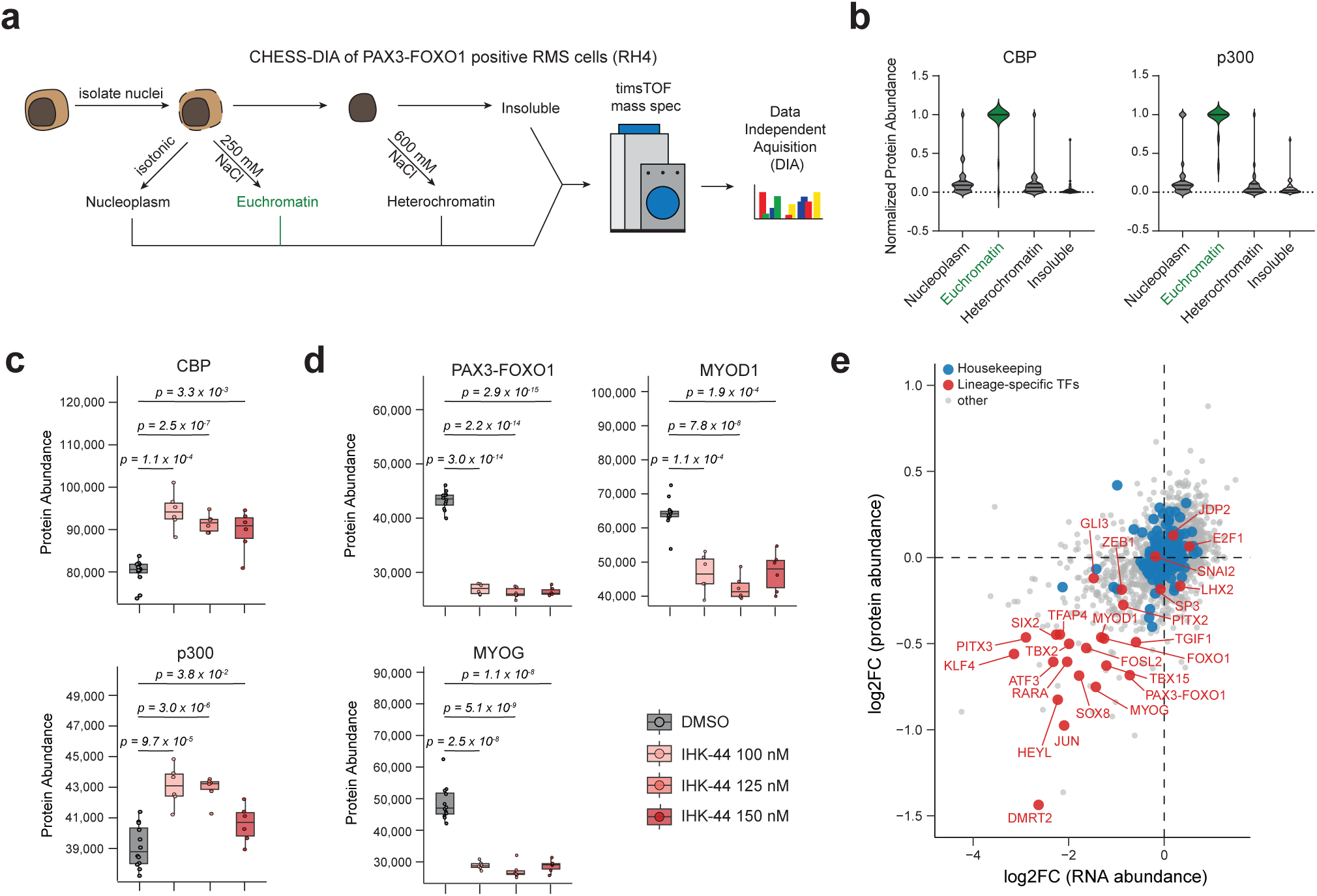
Chromatin focused mass-spec after CBP/p300 inhibition. **a.** Schematic of CHESS-DIA methodology. **b.** Violin plots of normalized protein abundance levels of CBP (left) and p300 (right) across nuclear compartments. **c.** Box plots of protein abundance levels between CBP (top) and p300 (bottom) in euchromatin fractions from RH4 cells treated with DMSO or increasing concentrations of IHK-44 (100 nM, 125 nM, 150 nM). Box plots represent median and quartiles, whiskers representing 1.5 x IQR. Statistical significance determined using a two-tailed Welch’s T-test. **d.** Box plots of protein abundance levels between PAX3-FOXO1, MYOD1, AND MYOG in euchromatin fractions from RH4 cells treated with DMSO or increasing concentrations of IHK-44 (100 nM, 125 nM, 150 nM). Box plots represent median and quartiles, whiskers representing 1.5 x IQR. Statistical significance determined using a two-tailed Welch’s T-test. **e.** Scatter plot comparing changes between RNA and protein abundance in RH4 cells treated with DMSO or 100 nM IHK-44 for 6 h. Lineage-specific TFs are highlighted and labeled in red, housekeeping genes in blue, other genes in gray.

Downregulation at enhancer-mediated genes is known to translate to the proteome with a temporal delay, reflecting the combined effects of reduced mRNA synthesis and limited protein stability^32^. To directly relate RNA-level and protein-level responses to IHK-44, we compared relative fold changes in transcript and protein abundance. Treatment of RH4 cells with 100 nM IHK-44 for 6 hours revealed a strong concordance between gene classes (**Figure 4e**). HK genes were largely resistant to depletion at both the transcript and protein levels, while LD genes exhibited reductions across both levels. Together, these data indicate that IHK-44 selectively downregulates lineage-defining genes which propagates to the proteome, disproportionately affecting proteins encoded by genes dependent upon enhancer-rich clusters.

### IHK-44 selectively impairs cell viability and induces growth arrest

Cell proliferation and survival are tightly coupled to the transcriptional programs that sustain lineage identity and growth-related pathways^3,33^. Because these programs arise through the cooperative interactions among multiple regulatory elements, even modest reductions in their activity can disproportionally affect cell fitness^34^. This sensitivity has been observed with previous epigenetic probes in FP-RMS, which has led us to consider the hypothesis that even partial inhibition of CBP/p300 catalytic activity may produce pronounced effects on cell viability^13,17,30,35,36^.

To test this hypothesis, we first examined how IHK-44 impacts cell viability in FP-RMS, fusion-negative (FN-RMS), and normal myoblast cells (**Figure 5a**). IHK-44 markedly reduced viability in FP-RMS cells, whereas FN-RMS and myoblasts cells were largely unaffected. Consistent with this, we monitored cell proliferation over time and saw a significant suppression of FP-RMS proliferation upon IHK-44 treatment, with minimal impact on FN-RMS or myoblasts (**Figure 5b**). To further evaluate this dependency, we tested IHK-44 and six other CBP/p300 modulators across a broader panel of RMS cell lines, which included five FP-RMS cell lines (RH4,

**Figure 5.**
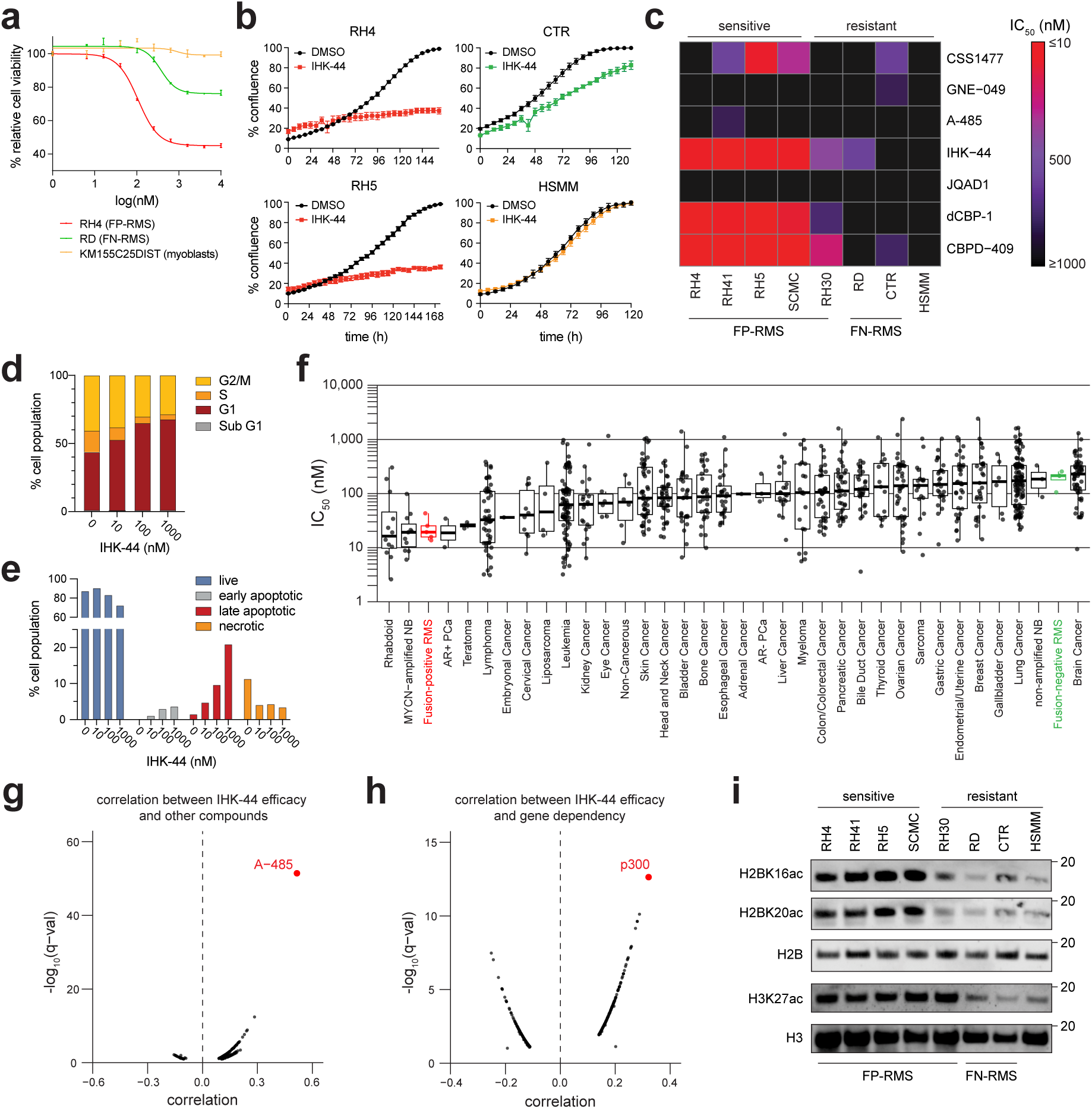
High H2B acetylation predicts antiproliferation response to CBP/p300 inhibition. **a.** Dose-response curves showing relative cell viability of FP-RMS (RH4), FN-RMS (RD), and normal myoblast (KM155C25DIST) cells after 72 h treatment with IHK-44. Values represent mean ± SEM of *n =* 4 technical replicates. **b.** Growth curves of % confluence over time (h) in FP-RMS (RH4, RH5), FN-RMS (CTR), and normal myoblast (HSMM) cell lines treated with DMSO or 185 nM IHK-44. Values represent mean ± SEM of *n =* 3 technical replicates. **c.** Heatmap summarizing IC_50_ values (nM) for CBP/p300 bromodomain inhibitors (CCS1477, GNE-049), CBP/p300 HAT domain inhibitors (A-485, IHK-44), p300 degrader (JQAD1), and CBP/p300 degraders (dCBP-1, CBPD-409) across sensitive and resistant RMS cell lines. **d.** Bar plots summarizing cell-cycle arrest of RH4 cells treated with increasing doses (10, 100, 1000 nM) of IHK-44 for 72 h. **e.** Bar plots summarizing apoptosis Annexin V/PI staining of RH4 cells treated with increasing doses (10, 100, 1000 nM) of IHK-44 for 72 h. **f.** IC_50_ values across 958 cell lines treated with IHK-44 from the PRISM screen grouped by primary disease. Box plot represents median and quartiles, whiskers showing 1.5 × IQR. Fusion-positive RMS is highlighted and labeled in red. Fusion-negative RMS in highlighted and labeled in red. **g.** Correlation between IHK-44 efficacy (log2.AUC) and other compounds in the DepMap Repurposing dataset. A-485 is highlighted and labeled in red. **h.** Correlation between IHK-44 efficacy (log2.AUC) and gene dependency scores in the DepMap CHRONOS dataset. p300 is highlighted and labeled in red. **i.** Western blots of non-treated histone acetylation levels in a panel of sensitive and resistant RMS cell lines. H2B and H3 are shown as loading controls

RH41, RH5, SCMC, and RH30), two FN-RMS cell lines (RD and CTR), and one normal myoblast cell line (HSMM). Across this panel, we saw that FP-RMS cell lines were highly sensitivity to IHK-44 with IC_50_ values less than 50 nM, while FN-RMS and myoblast cell lines were largely resistant with IC_50_ values greater than 500 nM (**Figure 5c and Supplemental Figure S6**). The same subtype sensitivity was also observed with the CBP/p300 degraders dCBP-1 and CBPD-409, consistent with previous Western blot results demonstrating rapid reductions in histone acetylation levels (**Supplemental Figure 2**). Interestingly, CBP/p300 bromodomain inhibitor CCS1477 was partially effective in some FP-RMS cell lines, while CBP/p300 inhibitor A-485, p300 degrader JQAD1, and CBP/p300 bromodomain inhibitor GNE-049 were largely ineffective.

We next investigated whether the antiproliferative effects of IHK-44 were driven from an arrest in the cell cycle, activation of apoptosis, or both. RH4 cells were treated with increasing concentrations of IHK-44 for 72 hours, leading to a pronounced G1 arrest with fewer cells in the S and G2/M phase (**Figure 5d and Supplemental Figure S7a**). Moreover, a slight increase in the number of apoptotic cells was observed at higher concentrations of IHK-44 (**Figure 5e and Supplemental Figure S7b**). Together, these findings indicate that IHK-44 selectively impairs FP-RMS proliferation primarily through cell cycle arrest, with apoptosis contributing at higher doses.

### H2B hyperacetylation predicts antiproliferative response to CBP/p300 inhibition

Motivated by the discovery that IHK-44 selectively suppresses FP-RMS proliferation, we next sought to determine if this selectivity extended to other cancer contexts and if there were molecular features that predicted sensitivity to CBP/p300 inhibition. To do so, we analyzed IHK-44’s activity in the Broad Institute’s Profiling Relative Inhibition Simultaneously in Mixtures (PRISM) 5-day cell viability screen across 958 cancer cell lines^8^. Results from the PRISM screen indicated that 431 cell lines (44.9%) had an IC_50_ of 100 nM or less while 21 cell lines (2.1%) had an IC_50_ of 1 µM or more (**Figure 5f**). As expected, FP-RMS cell lines were more sensitive than FN-RMS cell lines (FP-RMS IC_50_ = 23.4 nM; FN-RMS IC_50_ = 196.8 nM) (**Supplemental Figures S8a and S8b**). Among the lineages profiled, androgen receptor-positive prostate cancer (AR+ PCa) and MYCN-amplified neuroblastoma (MYCN-amp NB) models emerged as notably sensitive to IHK-44, whereas androgen receptor-negative (AR-) PCa and non-MYCN-amplified (non-amp) NB models were mostly resistant (**Supplemental Figures S8a and S8b**). Previous studies have shown that AR+ PCa and MYCN-amp NB are highly dependent on CBP/p300, and that CBP/p300 perturbation both dismantles the enhancer architecture that drives the disease and impairs cell viability^26,37^.

IHK-44 showed strong correlation with the CBP/p300 HAT domain inhibitor A-485 (**Figure 5g**). Bromodomain inhibitors (BRDi) were also correlated with IHK-44 response, consistent with previous findings that BRDi can serve as a therapeutic strategy in FP-RMS and other enhancer-addicted cancers (**Supplemental Figure S8c**)^30,38^. At the genetic level, IHK-44 response correlated with p300 gene dependency (**Figure 5h**). Members of the Spt-Ada-Gcn5 (SAGA) histone acetyltransferase complex (TADA1, TAFL5, TADA2B, TAF6L, and SUPT20H) and the Mediator complex (MED13L, MED24, MED19, MED23, MED13, and MED12) were also highly correlated, suggesting that IHK-44 preferentially affect cancers with a heightened demand for co-activator machinery (**Supplemental Figure S8d**)^39,40^.

The strong correlation between IHK-44 efficacy and co-activator dependency raised the possibility that cancers most sensitive to CBP/p300 inhibition share a common epigenetic framework. Cancers driven by enhancer-mediated transcription are often characterized by dense co-activator occupancy and high expression levels of LD genes, features that manifest from widespread hyperacetylation on histone tails^41,42^. Prior studies in AR+ PCa and MYCN-amp NB cell line models have shown that these enhancer-addicted states are marked by elevated H2B acetylation levels, a post-translational modification closely linked CBP/p300 catalytic activity^22^. Consistent with this, CBP/p300 inhibition by IHK-44 revealed AR+ PCa and MYCN-amp NB to be more sensitive than their counter-subtypes, AR-PCa and non-amp NB, respectively (**Supplemental Figures S8a and S8b**). These observations suggested that H2B hyperacetylation might represent a general epigenetic signature of enhancer dependency and could help explain why certain cancer subtypes display heightened sensitivity to CBP/p300 inhibition.

To test whether this principle also explains the subtype-selective sensitivity across FP-RMS and FN-RMS, we examined baseline histone acetylation levels across a panel of RMS cell line models. Notably, FP-RMS cell line models exhibited markedly higher levels of H2BK16ac and H2BK20ac levels while FN-RMS and non-RMS cell line models showed substantially reduced levels at these sites (**Figure 5i**). Together, these findings unite observations across enhancer-driven cancers to indicate that H2B hyperacetylation may serve as a potential biomarker of enhancer-addiction and vulnerability to CBP/p300 inhibition.

### CBP/p300 inhibition induces major RNA Pol2 loss at enhancer-regulated genes

Inhibition of RNA Pol2 dependent transcription can occur through a variety of mechanisms, as the life cycle of Pol2 moves through multiple sequential steps, primarily initiation, pausing, elongation, and termination. To determine the failure mode(s) of Pol2 resulting in downregulation of CBP/p300 dependent genes, we performed ChIP-seq of Pol2 after 6 hours of treatment with IHK-44. Then, we quantified changes in Pol2 across distinct sections of the gene body. These changes were then classified into 8 possible categories: promoter/TSSR ratio (increase: stalling, decrease: entry), TSSR/gene body ratio (increase: pause, decrease: release), gene body/TESR (increase: clogging, decrease: unloading), and total Pol2 gain or loss without shifts in gene body ratios (**Figure 6a**).

**Figure 6.**
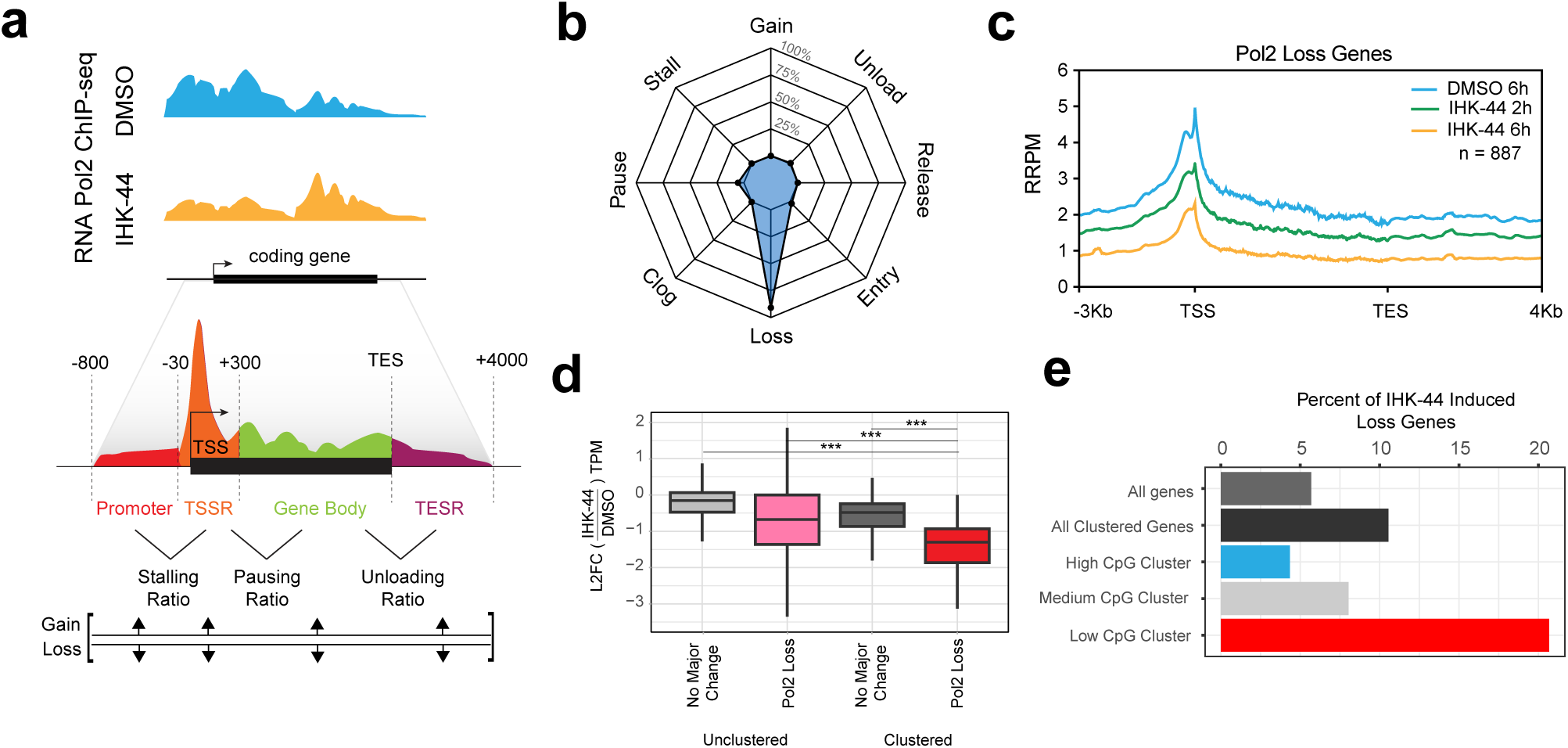
RNA Pol2 landscape changes stratify gene response to CBP/p300 inhibition. **a.** Schematic of the compPASS algorithm developed for the classification of RNA Pol2 alterations in response to drug treatment. **b.** Radar plot for the frequency of Pol2 changes in response to IHK-44 treatment for 6 hours in RH4 cells, indicating total Pol2 loss is most common, followed by Pol2 pausing for a smaller set of genes. **c.** Profile plots of Pol2 ChIP-seq signal across all genes in Pol2 loss category (*n = 887*), show loss between DMSO and IHK-44 treatment for 2 and 6 hours. **d.** L2FC TPM following IHK-44 treatment between weakly impacted (‘no major change”) and Pol2 loss genes, split between unclustered and clustered genes (box plots of median and quartiles, whiskers showing 1.5 × inter-quartile ranges; Mann-Whitney U test for significance) **e.** Percent of genes in various cluster categories that undergo total Pol2 loss after IHK-44 treatment for 6 hours.

Initial quantification of Pol2 across genes showed that 34% of genes had trivial amounts of Pol2 and thus were not included in further analysis. Of the rest, vast majority experienced no major change and only 4% of the total considered genes experienced a significant perturbation (**Supplementary Figure 9a**). Looking at just the perturbed set, analysis revealed that the dominant mode of CBP/p300i activity was to cause Pol2 loss across the entire gene body, with over >90% showing loss and few genes pausing (**Figure 6b**). This is in contrast to our previous results showing full degradation of CBP/p300 did not induce Pol2 loss, but rather Pol2 clogging (increase in the gene body, decreasing after the TES)^17^. This suggests non-catalytic scaffolding functions of CBP/p300 play an additional role in Pol2 distribution. Genes experiencing loss showed a strong trend of Pol2 depletion across the entire gene-proximal region following IHK-44 treatment for both two and six hours (**Figure 6c**). As an example, *MYC* shows loss in both conditions, with more severe Pol2 loss at six hours (**Supplementary Figure 9b**).

Next, we tested a dual hypothesis: (1) that genes in the Pol2 Loss category were more downregulated at the mRNA level, and that (2) genes in the Pol2 Loss category were in clusters with low CpG island content. Splitting genes into groups according to Pol2 Loss and cluster status indeed showed both that genes with Pol2 loss inside clusters were the most strongly downregulated (**Figure 6d**). Moreover, CpG island deficient clusters contained the greatest number of genes losing Pol2 (>20%), while clusters with high frequency of CpG islands had the fewest number of genes undergoing Pol2 loss (<5%) (**Figure 6e**). This trend holds when looking at the log2 fold-change in total Pol2 across all genes, with increasing CpG island frequency correlating with increased Pol2 loss (**Supplementary Figure 9c**). Given that more enhancer-driven genes tend to occupy low CpG island content clusters, CBP/p300i appears to preferentially deplete Pol2 at LD genes rather than induce CpG-mediated Pol2 accumulation.

### Machine learning identifies gene and cluster features predictive of CBP/p300 dependence

The degree to which any given gene depends on the acetylating activity of CBP/p300 likely depends on a combination of multiple gene features. Building on previous observations, we hypothesized that architectural enhancer features (the extent of 3D enhancer clustering and Pol2 connectivity) combine with RNA transcript stability (mRNA half-life) to define the genes most acutely dependent on CBP/p300 activity. For relative mRNA stability^43^, we measured nascent RNA with ChRO-seq and estimated the half-life of transcripts as the ratio of steady-state levels (RNA-seq) to RNA production levels (ChRO-seq) (**Supplementary Fig 10a**). As shown in prior work, the half-life metric segregates HK genes from LD genes, as exemplified by distinct 2D and half-life rank positions of *GAPDH* and *MYC* (**Supplementary Figures 10b and 10c**).

We then built a Random Forest model using groups of features, including cluster composition, RNA Pol2 occupancy, TF occupancy, CpG island abundance, and relative mRNA half-life, to predict each gene’s response to CBP/p300 inhibition (IHK-44) (**Figure 7a**). First, we used a classifier to predict whether a gene is downregulated (L2FC from IHK-44 < 0.999). We trained models for predefined subgroups of features and all features over 100 random splits and measured performance. Classification using all features (shown in red) performed best with high precision, high recall and a strong receiver operating characteristic (ROC) curve (**Figures 7b and 7c**). When using the same features but a random forest regression, using all features had the least predictive error and highest median R^2^ (**Supplementary Figures 10d and 10e**).

**Figure 7.**
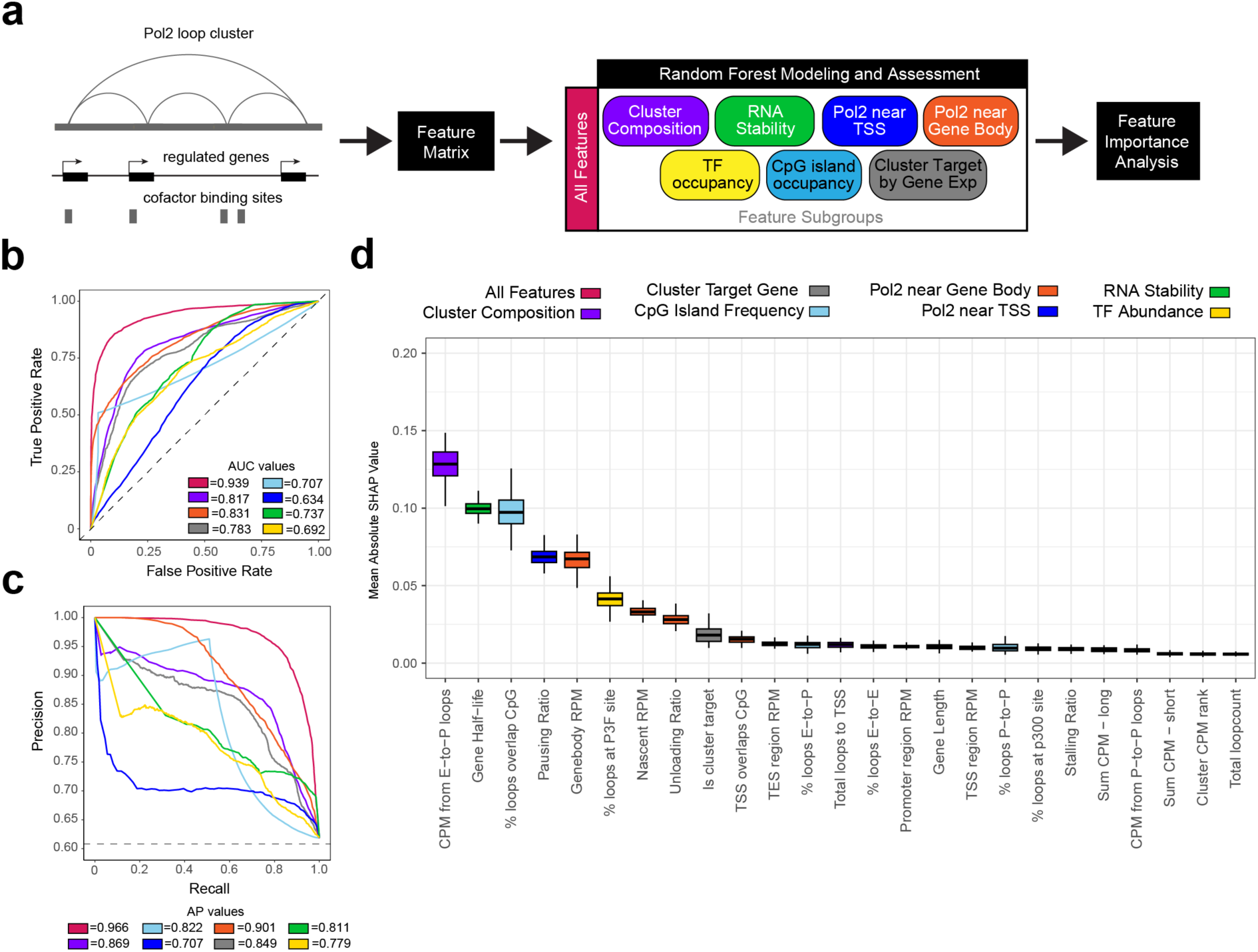
Identification of gene and cluster features that predict transcriptional dependence on CBP/p300. **a.** Design of a Random Forest machine learning model for predicting response to CBP/p300 inhibition at 6 hours, considering a mix of gene features, followed by SHAP permutation to learn the relative importance of each feature to predict responsiveness. **b.** The ROC (true positive versus false positive) curves for classification models from each gene feature group, and the combination of all features. Each line is colored according to the feature colors in (**a.**). AUC, area under the curve. **c.** The precision-recall curves for classification models from each gene feature group, and the combined all feature performance. Each line is colored according to the feature colors in (**a.**). AP, Average Precision. **d.** SHAP feature importance analysis colored by feature group across 100 regression runs. Each boxplot is colored according to the feature group colors in (**a.**).

As these feature groups could conflate multiple individual features, we next performed a SHapley Additive exPlanations (SHAP) test on the All Features Random Forest regression. Taking the mean absolute SHAP values for each condition across 100 models, we ranked the importance for each feature, colored by subgrouping (**Figure 7d**). The results show that cluster strength (as measured by total CPM at enhancer-promoter loops), RNA half-life and percent CpG-islands in loop anchors are the most important features for predicting responsiveness to CBP/p300 inhibition. Interestingly, many individual features can be completely discarded without suffering a loss in predicting gene responses to IHK-44, including gene length, the overall number of loops in a cluster, the presence or absence of a CpG island in the target gene’s promoter, or the percentage of enhancer-enhancer/enhancer-promoter/promoter-promoter loops in the cluster. The high accuracy of the classification model with all features supports the concept that predictive power depends on training a model with a high diversity of these critical gene and cluster features.

## Discussion

Our findings herein define CpG island-dependent chromatin as a central organizing principle of Pol2-mediated clustering. Within this framework, we identified two distinct classes of Pol2-mediated clusters: one characterized by high CpG island content, P-P interactions, and expression of HK genes, and another defined by high CBP/p300 occupancy, E-P looping, and expression of LD genes. Acute inhibition of CBP/p300 catalytic activity results in the rapid loss of acetylation at enhancers and preferential downregulation of LD transcriptional networks, leading to impaired cellular proliferation and activation of apoptotic programs. We also demonstrated that cluster strength, RNA half-life, and CpG island content are the most important features for predicting CBP/p300 inhibition. Collectively, our findings establish a model in which CpG island-associated chromatin governs Pol2-mediated cluster organization and determines gene-specific sensitivity to CBP/p300 inhibition.

CpG island content strongly stratified cluster architecture and predicted sensitivity to CBP/p300 inhibition. Several studies support a mechanistically instructive role for CpG islands. CpG-rich DNA sequences are generally unmethylated and inherently adapted for promoter function, destabilizing nucleosomes, and attracting chromatin regulators to create a transcriptionally permissive state^16^. As a proof-of-concept, the introduction of an artificial CpG island into mouse cells has been shown to induce epigenetic features that are characteristic of promoters, suggesting that CpG islands act as promoters by default^44^. The strong and consistent association between CpG island content, promoter-centric Pol2 clustering, and transcriptional resilience to CBP/p300 inhibition suggests that CpG islands, when arrayed closely in a manner than enables clustering, collectively maintain HK gene expression. It is possible that nearby CpG-rich promoters act as pseudo-enhancers to nearby genes. It has been shown that the promoters of lncRNAs can act as “enhancers” to nearby genes^45^. While CpG island presence or absence does not predict gene expression, or responsiveness to CBP/p300 inhibition, dense arrays of 3D connected CpG islands does.

Our findings integrate with a growing body of work describing enhancer-driven transcriptional addiction and dependence on CBP/p300 in cancer. Prior studies have shown that tumors reliant on cell-identity defining transcriptional programs frequently harbor dense enhancer assemblies at lineage-defining genes that require high levels of acetylation to maintain transcriptional output^17,22,26,30,41^. Our results extend this framework by linking enhancer dependency to Pol2 cluster architecture and by showing that CpG island content separates stable promoter-centric loci from enhancer-driven, acetylation-dependent loci. In this context, H2B hyperacetylation has emerged as a candidate marker of enhancer dependency, potentially reflecting the rapid acetylation kinetics required to sustain transcription at lineage-defining genes^21,22^. Prospective validation of enhancer-associated H2B acetylation in patient-derived samples will be required to determine its utility as a predictive biomarker in clinical settings.

In summary, our study connects DNA sequence composition, 3D chromatin architecture, and enhancer addiction to define a mechanistically grounded framework for targeting CBP/p300-dependent transcriptional programs.

### Limitations of study

A key limitation of this study is that our model was derived from the RH4 cell line, and the extent to which CpG island-dependent chromatin and Pol2 organization generalizes across diverse lineages, cellular differentiation states, or non-malignant tissues remains to be determined. The classification of LD genes is respective to our model system, and refinement of these categories across broader biological contexts may further clarify how enhancer-driven LD genes are organized. Our study was designed to specifically examine and pharmacologically perturb features or regulators distinguishing Pol2-associated chromatin interactions, and therefore does not address how other DNA sequence features, transcriptional regulators, or chromatin-associated complexes may contribute to higher-order genome organization.

## Resource availability Lead contact

Further information should be directed to and will be fulfilled by the lead contact, Berkley E. Gryder.

## Materials availability

Synthesis details for IHK-44 are provided in the Supplemental Information and are available from the lead contact with a completed materials transfer agreement.

## Data and code availability

Results from the PRISM screening are provided in the Supplementary Tables (Table S1-S3). All next-generation sequencing datasets (RNA-seq, ChIP-seq, ChRO-seq) have been deposited in the NCBI’s Gene Expression Omnibus under accession number GSE319181. Mass spectrometry proteomics datasets (CHESS-DIA) will be deposited on Pride. Original Western blot images will be deposited on Mendeley Data. This study does not report original code. Scripts used throughout this study are available at https://github.com/GryderArt and https://github.com/mattychang. Any additional information required to reanalyze the data reported in this study is available from the lead contact upon request.

## Acknowledgements

We would like to thank the Case Western Reserve University School of Medicine’s Genomics Core, especially Simone Edelheit, for their assistance with the next-generation sequencing experiments. We would like to thank the Case Comprehensive Cancer Center’s Cytometry and Imaging Microscopy Shared Resource (P30CA043703) for assistance with the flow cytometry experiments. We would like to thank the PRISM team at the Broad Institute of MIT and Harvard for conducting the PRISM screen. B.E.G. was supported by the NCI of the National Institute of Health under award number 1R01CA291963-01, DOD’s Convergent Science Virtual Cancer Center under award number W81XWH-21-1-0298, Alex’s Lemonade Stand Foundation Crazy 8 “A” award, and B+ foundation. D.H.C. was supported by the NIGMS of the National Institutes of Health under award numbers T32GM007250 and T32GM135081, as well as by the NCI under award number F30CA294663.

## Author contributions

**M.I.H. Khan**: Conceptualization, data curation, formal analysis, supervision, validation, investigation, visualization, methodology, project administration, writing–original draft, writing–review and editing. **M.S. Chang**: Conceptualization, investigation, data curation, formal analysis, validation, investigation, visualization, methodology, writing–original draft, writing–review and editing. **Y. Asante**: Formal analysis, data curation, visualization, writing–original draft. **T. Kukhta**: Formal analysis, data curation, visualization. **M.K. Al-Haddad**: Methodology, investigation, formal analysis, validation. **B. Udhayakumar**: Methodology, investigation, formal analysis, validation. **A.M. Garzón-Porras:** Investigation, validation. **E.L. Rotariu**: Investigation, validation, data curation. **S. Aneja:** Formal analysis, data curation. **J.L. Kelly**: Investigation, validation. **C. Farma-Hai**: Investigation, validation. **K. Ullal:** Formal analysis, data curation. **D.H. Chin**: Investigation, data curation, funding acquisition. **A.J. Federation**: Methodology, investigation, validation, formal analysis. **L.K. Pino**: Methodology, investigation, validation, formal analysis. **J. Robbins**: Methodology, investigation, validation, formal analysis. **M. Wachtel**: Investigation, resources, data curation, formal analysis, writing–review and editing. **S.R. Stauffer:** Resources, writing-review and editing. **B.E. Gryder**: Conceptualization, resources, data curation, formal analysis, supervision, funding acquisition, validation, investigation, visualization, methodology, writing–original draft, project administration, writing–review and editing

## Methods

### Chemicals

All small molecules were dissolved in DMSO and stored at concentrations of 10 mmol/L at −20°C or −80°C. All small molecules were diluted to a final concentration of ≤0.3% by volume in media for all cell culture experiments. Synthesis details for IHK-44 are provided in the Supplementary Methods. A-485 and GNE-049 were purchased from Selleck Chem (cat. #S8740, cat. #S8625). CCS1477, JQAD1, dCBP-1 were given as a gift from Jun Qi (Dana-Farber Cancer Institute, Cambridge, MA). CBPD-409 was given as a gift from Shaomeng Wang (University of Michigan Medical School, Ann Arbor, MI).

### Cell Culture

FP-RMS cell lines, RH4, RH41, RH30, and SCMC, were cultured in DMEM (Gibco, cat. #10313039) supplemented with 10% fetal bovine serum (FBS; Gibco, cat. #A3160402) and 1x penicillin-streptomycin-glutamine (Gibco, cat. #10378016). Luciferase reporter cell line (RH4-mALK), FN-RMS cell lines (RD and CTR), and mouse myoblast C2C12 cells (ATCC, cat. #CRL-1772), were grown in the same media as FP-RMS cells. FP-RMS cell line RH5 and immortalized human skeletal muscle myoblasts HSMM (provided by Heide Ford, University of Colorado School of Medicine, Aurora, CO, USA with permission from Corinne M. Linardic, Duke University, Durham, NC, USA) were maintained in SkGM-2 media bullet kit (Lonza, cat. #CC-3245). All cells were cultured in 5% CO2 at 37°C. Cell lines were authenticated upon receipt by short tandem repeat profiling and used for experimentation from frozen stocks. All cells were regularly tested for mycoplasma contamination using the MycoStrip™ assay (InvivoGen, cat. #rep-mysnc-50).

### HiChIP Loop and Cluster Calling

Clusters were identified from HiChIP data via the Genic Rank of Active Clustered Elements (GRACE) pipeline. Briefly, we used MACS3 to call peaks from RNA Pol2 HiChIP data in a linear format. We then built all possible interactions between peaks with possible interactions restricted by pre-defined topologically associating domains (TADs) and distance thresholds (minimum 8 kb, maximum 3 Mb). We then annotated loops with contacts per million (CPM) values from a corresponding contact matrix at a 5 kb resolution. We removed zero CPM loops and carried out a quantile regression (per chromosome) to capture the top 85^th^ percentile of loops. Using the remaining filtered loops, we called clusters as any set of loops with overlapping anchors that daisy chained to at least 3 loops. We assigned cluster-associated genes per cluster by selecting all genes where the TSS is within 5 kb +/- of a loop anchor. We used RNA-seq to determine the most expressed gene of each cluster and assigned that as the target gene (for each cluster which contained genes).

### Molecular Docking

In silico molecular docking studies were performed using the Schrödinger Suite 2024-4 (Schrödinger, LLC, New York, NY). The docking protocol involved three major steps: (a) ligand preparation, (b) receptor grid generation, and (c) molecular docking.

### Ligand Preparation

Ligands were prepared using the LigPrep tool (Schrödinger, 2024-4). The 2D structures were imported in SDF format. The OPLS4 force field was used for energy minimization. Possible ionization states were generated at pH 7.0 ± 2.0 using Epik Classic, and the specified chirality was retained.

### Protein Preparation and Grid Generation

The protein crystal structure was imported from the Protein Data Bank (PDB ID: 8GZC) and prepared using the Protein Preparation Workflow wizard. The workflow includes four steps: specify protein, preprocess, optimize H-bond assignments, and minimize and delete waters. Bond orders were assigned, hydrogens were added, missing side chains and loops were filled using Prime, and water molecules beyond 5.0 Å from any hetero group were removed. Hydrogen bond assignments were optimized using the PROPKA. The waters were removed from the ligands (hets), and the structure was minimized using the OPLS4 force field. A receptor grid was generated using the Receptor Grid Generation panel. The active site was defined using the co-crystallized ligand. Van der Waals scaling factors of 1.0 and a partial charge cutoff of 0.25 were applied to soften nonpolar receptor atoms. The enclosing box was prepared with the centroid of the workspace ligand as its center, and the dock ligand’s length was modified using advanced settings.

### Ligand Docking

Prepared ligands were docked into the receptor grid using the Ligand Docking module in Standard Precision (SP) mode. Default parameters were used unless otherwise specified. The top poses were selected based on the docking scores, and binding interactions were further analyzed using Maestro’s Pose Viewer panel.

### Western Blotting

RH4 cells were plated at 10,000,000 cells in 15 cm dishes. After 24 hours of seeding, cells were treated at the indicated concentration and time and collected by trypsinization. For all experiments, histones were extracted using the histone extraction kit (Abcam, cat. #ab113476) and quantified using a BCA assay (Bio-Rad, cat. #5000006). Histones were separated using a 4-12% Bis-Tris SDS-PAGE gel (Invitrogen, cat. #NP0321BOX; 60 V for 15 min, 120 V for 120 min) and transferred to a nitrocellulose membrane (Amersham Protran, cat. #10600015; 200 mA for 120 min). The membrane was blocked in 5% milk in TBS/0.1% Tween for 60 minutes at room temperature and incubated with primary antibodies in 2.5% milk in TBS/0.1% Tween overnight at 4°C. After washing the membrane three times with TBS/0.1% Tween, the membrane was incubated in Precision Protein™ StrepTactin-HRP Conjugate (Bio-Rad, cat. #1610380) and HRP-conjugated secondary antibody for one hour. Three additional washes in TBS/0.1% Tween and one wash in TBS were completed before visualization. Histones were detected by chemiluminescence using an ECL detection reagent (Thermo Fisher cat. #34094). Images were captured using the BioRad ChemiDoc Imaging System and analyzed with the Image Lab 6.1.0 software. The following antibodies were used: anti-H2BK16ac (1:2000, Abcam, cat. #ab177427), anti-H2BK20ac (1:2000, Abcam, cat. #ab177430), anti-H2B (1:1000, Active Motif, cat. #39210), anti-H3K27ac (1:2000, Active Motif, cat. #39133), anti-H3 (1:2000, Abcam, cat. #ab1791), and anti-rabbit IgG (1:5000, Cytiva, cat. #45-000-682).

### ChIP-seq

#### Sample Preparation

RH4 cells were plated at 10,000,000 cells per well into 15-cm dishes. After 24 hours of seeding, cells were treated with IHK-44 at 100 nM for 30 minutes, 2 hours, or 6 hours, or 6 hours with DMSO. At the indicated time points, cells were collected by trypsinization. Cells were fixed with 1% paraformaldehyde, quenched with 2.5 M glycine, and snap-frozen at −80°C overnight. DNA was digested using a MNase reaction mix (10 µL of 10X Micrococcal Nuclease Buffer (New England Biolabs, cat. #B0247SVIAL), 0.5 µL of 200X Recombinant Albumin (New England Biolabs, cat. #B9200SVIAL), 1.30 µL of Micrococcal Nuclease (New England Biolabs, cat. #M0247SVIAL), and 89.5 µL of dH_2_O). After incubation in a shaker for 25 minutes at 25°C, the digestion reaction was stopped with 10 µL of 0.5 M EGTA. Cells were lysed using a RIPA buffer (20 µL of 10X RIPA (Dovetail HiChIP *MNase* Kit v2.1), 8 µL of protease inhibitors (Roche, cat. #11836145001), 2 µL of HALT protease inhibitors, (Thermo, cat. #1861284), and 60 µL of dH_2_O). After incubation on ice for 30 minutes, lysates were centrifuged at 20,000 xg at 4°C for 15 minutes. For histone acetylation markers, lysates were spiked with chromatin extracted from mouse myoblast C2C12 cells at a defined and conserved ratio of 1:20 C2C12 chromatin to human chromatin and incubated with a conserved ratio of 1:1 antibody to human chromatin overnight (anti-H2BK16ac, Abcam, cat. #ab177427; anti-H2BK20ac, Abcam, cat. #ab177430). Lysates were then immunoprecipitated with Protein A/G beads (Thermo, cat. #78609), washed 4x with a low-salt RIPA wash buffer (50 mM Tris-HCl, 1% NP40, 0.1% SDS, 150 mM NaCl), 2x with a LiCl wash buffer (10 mM Tris-HCl, 250 mM LiCl, 1% NP40, 1% NaDoc, 1 mM EDTA), 2x with a TE buffer (10 mM Tris-HCl, 1 mM of EDTA), de-crosslinked with 17.5 µL of PK buffer (50 mM Tris-HCl, 25 mM EDTA, 1.25% SDS) and 6 µL of Proteinase K (Sigma-Aldrich, cat. #3115828001), followed by DNA purification (Zymo Research, cat. #D4013). For RNA Pol2 (anti-Pol2, Santa Cruz Biotechnology, cat. #sc-47701 X), lysates were incubated overnight with Dynabeads (Thermo, cat. #14311D), washed 2x with a high-salt RIPA wash buffer (50 mM Tris-HCl, 1% NP40, 0.1% SDS, 500 mM NaCl), 2x with a low-salt RIPA wash buffer, 2x with a TE wash buffer, de-crosslinked, and purified as described before. Libraries were prepped accordingly (New England Biolabs, cat. #E6440S and #E7645S), and quantity and fragment size were measured using a Qubit 4 fluorometer (Thermo Fisher) and a TapeStation (Agilent). Sequencing was performed on Illumina NextSeq 550 High Output Flowcell.

#### Data Processing and Analysis

ChIP-seq FASTQ files were aligned to the human genome hg38 and spike-in mouse genome mm10 using BWA (v0.7.17). Reads-per-million (RPM) values were quantified after fragment extension and shifting corrections. Peaks were called using MACS3 (v3.0.0a6) and HOMER (v4.9.1). Regions that were called as peaks but are overlapping black-listed regions defined by the ENCODE consortium were removed before further analysis. Heatmaps and metagene plots were generated with a custom plotBEDHeat.sh pipeline that implements the bedtools and deeptools packages. For multi-sample analysis, we used a custom runBedCovCompSpiked.sh pipeline that reads from multiple BAM files to generate normalized densities at BED peak regions. Previously determined RPM values were normalized using the mouse spike-in, where relative RPM (RRPM) values are defined by the following equation: RRPM = (human RPM in peak x 10^6^) / total mouse reads. Super enhancers were identified using the ROSE2 package with a stitching parameter of 12,500 bp. Genome-browser visualization was performed by using IGV. Visualization of multi-sample analysis and super enhancers were performed by using custom R scripts (https://github.com/GryderArt/ChIPseqPipe).

#### Luciferase Assays

Luciferase reporter cell line, RH4-mALK, was plated 2,000/well in white-walled, white-bottom 384-well white-walled, white-bottom plates (Thermo, cat. #164610). 24 hours after seeding, cells were dosed using a range of 8 concentrations starting from 10µM (dilutions divided by 3) and were performed in triplicate. 24 hours after treatment, luminescence was detected using the Bright-Glo™ Luciferase Assay System (Promega, cat. #E2620) on an automated plate reader (Thermo Varioskan LUX). IC_50_ values were calculated for each time point using GraphPad Prism (v10.5.0) statistical analysis for non-linear regression. Details on the generation of RH4-mALK and RH4-CMV cell lines are as previously described^35^.

### RNA-seq

#### Sample Preparation

RH4 cells were plated at 500,000 cells per well into 6-well plates. After 24 hours of seeding, cells were treated with A-485, dCBP-1, IHK-44, and JQAD1 at 100 nM and 1000 nM, or equal volumes of vehicle DMSO, for 2 hours or 6 hours, and then collected by trypsinization. Samples were lysed and homogenized using Qiashredder homogenizer (Qiagen, cat. #79654), and total RNA was extracted from the cells using RNeasy Plus Mini Kit (Qiagen, cat. #74136). Quantity was measured by a Qubit 4 fluorometer (Thermo Fisher). Libraries were prepped accordingly (New England Biolabs, cat. #E7765S). Sequencing was performed on Illumina NextSeq 550 High Output Flowcell.

#### Data Processing and Analysis

RNA-seq FASTQ files were aligned to the human genome hg38 using STAR (v2.5.3a). Transcript-per-million (TPM) values were quantified using RSEM (v1.3.3). Gene set enrichment between two samples was determined using the NCI’s Gene Set Enrichment Analysis (GSEA) tool (v4.3.3). Custom gene set references were created by selecting up- and down-regulated gene sets from the log_2_ fold change (log2FC) metric, followed by rank-ordering. Genome-browser visualization was performed using IGV. Visualization and summary of GSEA results were performed using custom R scripts (https://github.com/GryderArt/VisualizeRNAseq).

#### CHESS-DIA Proteomics

CHESS-DIA was performed as previously described^31^. Briefly, RH4 cells were plated at 200,000 cells per well in 96-well plates and treated in quadruplicate with 100 nM, 125 nM, or 150 nM of IHK-44, or an equivalent volume of DMSO. After 6 hours, the cells were collected, their nuclei extracted, and then fractionated on beads via KingFisher. The chromatin fraction was digested with the AutoSP3 protocol. After the digestion was quenched, the peptides were diluted to an injection concentration of 2 ng/µL in 0.1% FA and 100 ng was loaded onto EvoTips before being analyzed using diaPASEF on the timsTOF Ultra. A pool of samples was searched alongside the experimental injections to improve match between runs in DIA-NN (v1.8.1).

#### Cell Viability Assays

Three thousand cells in a volume of 20 μl were seeded per well in a 384-well plate (Greiner, 7.781 098). One day later, IKH-44 was added to the cells using an HP D300 digital dispenser (Tecan, Switzerland). After 3 days of treatment, cell viability was measured using the CellTiter-Glo 2.0 cell viability assay (Promega, G9242). Dose-response curves were fitted by nonlinear regression using GraphPad Prism 8.0.0.

#### Cell Proliferation Assays

RH4, RH5, RH30, RH41, SCMC, RD, CTR, and HSMM cells were plated in 384-well plates to achieve 15% confluence at time of drug dosing and monitored until control (DMSO) wells reached >95% confluence. 24 hours after seeding, cells were dosed using a range of 11 concentrations starting from 5µM (dilutions divided by 3) and were performed in triplicate. The dose-response was quantified by percent cell confluence from phase contrast images taken every 6 hours using the IncuCyte ZOOM software. IC_50_ values were calculated for each time point using GraphPad Prism (v10.5.0) statistical analysis for non-linear regression.

#### Flow Cytometry Cell cycle arrest

RH4 cells were plated at 250,000 cells per well into 6-well plates and treated with 10, 100, and 1000 nM, or equal volumes of DMSO, for 72 hours. Floating and adherent cells were collected and resuspended in 50 μl of PBS. 500 μl of precooled 70% EtOH (−20°C) was then added dropwise to the cells under gentle vortex. After incubation of the cells at −20°C overnight, cells were pelleted by centrifugation at 1000xg for 5 min and washed once with PBS. The cell pellet was then resuspended in 500 μl of a solution containing 20 mg/ml propidium iodide (Thermo, cat. #P1304MP), 200 μg/ml RNAse A (Thermo, cat. #EN0531) and 0.1% Triton X-100. Cells were then incubated in the dark for 15 min on ice and analyzed by flow cytometry (BD Biosciences LSRFortessa). Cell cycle distribution was analyzed by FlowJo (v11.0.2.158272).

#### Apoptosis

RH4 cells were plated at 250,000 cells per well into 6-well plates and treated with 10, 100, and 1000 nM, or equal volumes of DMSO, for 72 hours. Floating and adherent cells were collected and resuspended in 500 μl of binding buffer (Thermo, cat. #V13246), 2.5 μl of Annexin V-FITC (Thermo, cat. #A13199), and 2.5 μl of propidium iodide (Thermo, cat. #P1304MP). Cells were then incubated in the dark for 15 min at room temperature and analyzed by flow cytometry (BD Biosciences LSRFortessa). Apoptosis distribution was analyzed by FlowJo (v11.0.2.158272).

#### PRISM Screening

PRISM screening was performed as previously described^8^. Briefly, IHK-44 was screened in a 958 DNA-barcoded cell line pool, where 20-25 cell lines per pool were plated in 384-well plates and treated with IHK-44 in a range of 8 concentrations starting from 3 µM (dilutions divided by 3) in triplicate for 5 days. Cells were lysed with DNA lysis buffer, DNA was extracted via KingFisher, and PRISM barcodes were amplified by PCR. Detection of the barcodes and analysis was then performed as previously described.

#### Characterization of RNA Pol2 Landscape

Gene loci are segmented into four regions relative to transcription start and end sites: promoter (−800 to −30), TSS (−30 to +300), gene body (+300 to TES), and TES (TES to +4000). Genes shorter than 2 kb and genes on chromosome Y were excluded. We used bedtools (v2.30.0) to quantify Pol2 ChIP-seq reads within these four regions. To avoid quantifying trivial Pol2 values, we remove genes with low promoter-proximal Pol2 signal (bottom 20th percentile dropped). Normalized read densities (RPM per base) are computed for each region and pausing, stalling, and unloading ratios are defined as the density of an upstream region divided by that of the downstream region. Genes are classified by comparing case (drug-treated) versus control log2 fold changes in regional signal and ratios using thresholds predefined from biological replicates of Pol2 ChIP-seq data. Final states for each gene (e.g., pausing vs releasing) are assigned based on the ratio with the greatest magnitude and its sign, allowing multiple candidate states but enforcing a single final classification per gene. Genes not assigned to any of the primary classes (stalling, pausing, clogging, entering, releasing and unloading) may be assigned to the state “Pol2 gain”, if the log2(Prom + TSS + Gene Body + TES reads from case) – log2(Prom + TSS + Gene Body + TES reads from control) > 0.999 or “Pol2 loss” is the inverse condition of gain is true. Genes which meet none of the criteria are marked as “no major change”. Signal at identified gene sets was visualized with deeptools (v3.5.5).

### Random Forest Modeling

#### Dataset Generation and Feature Subgrouping

A total of 25 epigenetic- and transcriptomic-based features were obtained from RNA-seq, RNA Pol2 AQuA-HiChIP, RNA Pol2 ChIP-seq and ChRO-seq analyses. For all genes overlapping the span of a HiChIP cluster, gene response to IHK-44 was expressed as both continuous and binary values. For the former, the L2FC of the transcripts per million (TPM) in the presence of IHK-44-treated and DMSO-treated cells was measured. For the latter, genes with an L2FC < −0.999 were labeled as the positive class (downregulated), while the remaining genes were labeled as the negative class (not downregulated). The result was a dataset with 3,809 total observations which, in the case of the binary response variable, had 552 cases of downregulated genes. Features were grouped into 7 subgroups based on both their correlation within the dataset and practically based on their source. The assignment of features to subsets and variable types for the corresponding features are annotated in Supplemental Table S4.

#### Model Training

We used Random Forest regression on the continuous response variable and classification on the binary response variable (n_estimators set to 100). To obtain a more robust measure of performance, five-fold cross-validation was performed using 20 different random splits, resulting in a total of 100 models being independently trained, with each sample being tested 20 times. Because the L2FC is a continuous variable, a random forest regression model was trained to relate features to the magnitude of differential expression. Separately, we trained a random forest classification model on the same dataset to focus on the most strongly downregulated genes. The same process that was applied to all features was applied to the 7 subgroups. Training was performed with the Scikit-Learn library (version 1.6.1) in Python.

#### Model Evaluation

For the regression models, we evaluated model performance by comparing mean-squared error (MSE) and the R² score distributions from all 100 trained models. For the classification models, we measured performance in terms of precision, recall, the average precision of the precision-recall curve (AP/AUCPR), and the area under the receiver operating characteristic curve (AUCROC) for all 100 trained models.

#### Feature Importance Analysis

The relative importance of each feature was computed using a computed average absolute SHAP values for each feature. SHAP importance was measured with the TreeExplainer function from the SHAP library (version 0.50.0) in Python.

#### ChRO-seq

Chromatin run-on sequencing (ChRO-seq) libraries were generated from RH4 rhabdomyosarcoma cells using a streamlined adaptation of the Danko laboratory protocol^46^. The protocol requires an RNase free environment thus SUPERase•In RNase inhibitor (Thermo Fisher Scientific, cat. AM2694) was used in the experiment reactions and working buffers while the equipment and work bench were cleaned with RNase-zap (Sigmaaldrich, cat. R2020-250ml). The chromatin was isolated via NUN lysis buffer, fragmented via vortexing and DNase I reaction (NEB DNase I cat. M0303L & 10x DNase I reaction buffer cat. B0303S) was subjected to nuclear run-on in the presence of biotin-11–labeled ribonucleotides (PerkinElmer/Revvity Biotin-11-ATP cat. NEL544001EA, Biotin-11-CTP cat. NEL542001EA, Biotin-11-GTP cat. NEL545001EA, Biotin-11-UTP cat. NEL543001EA) to label nascent RNA which resulted in various nascent RNA fragment sizes, followed by RNA extraction with TRI Reagent LS (MRCgene cat. TR-120 or Thermo Fisher Scientific cat. 10296-010) and phase separation using 1-bromo-3-chloropropane (BCP) (Sigma-Aldrich cat. B9673) to reduce DNA contamination. Biotinylated Nascent RNA was ethanol-precipitated with GlycoBlue coprecipitant (Thermo Fisher Scientific cat. AM9515), followed by base hydrolysis and buffer-exchanged with Micro Bio-Spin P-30 gel columns (Bio-Rad, cat 732-6250) prior to adaptor ligation. A 3′ RNA adaptor was ligated off-bead using T4 RNA ligase I and 10x T4 RNA ligase buffer (NEB cat. M0204L) in the presence of ATP, and PEG 8000, with adaptor concentration scaled to chromatin input to minimize adaptor dimers. Biotinylated RNA was first enriched using hydrophilic streptavidin magnetic beads (NEB, S1421S). Then, the 5’ end was decapped on-bead with RNA 5′ pyrophosphohydrolase RppH (NEB cat. M0356S) in Thermopol buffer (NEB cat. B9004S) and phosphorylated on-bead with T4 polynucleotide kinase (NEB cat. M0201L). Following a second streptavidin bead enrichment and a RNA extraction with TRI Reagent (MRCgene cat. TR-118 or Thermo Fisher Scientific cat. 15596026), BCP, and GlycoBlue, a 5′ RNA adaptor was ligated off-bead. After, a third streptavidin bead enrichment and RNA extraction is performed followed by RNA reverse transcription using SuperScript IV reverse transcriptase (Thermo Fisher Scientific, cat. 18090010) and clean-up. Libraries were amplified by PCR using indexed primers from IDT and SsoAdvanced Universal SYBR Green Supermix (Bio-Rad cat. 1725271), with cycle number determined empirically by qPCR, purified by column cleanup, assessed with Qubit fluorometry and TapeStation, followed by size selection using Pippin Prep machine and 2% agarose gel internal marker (Sage Science machine cat. PIP000, 2% gel, cat. CDF2010), and assessed again by Qubit fluorometry and TapeStation prior to sequencing. Sequencing reads from were processed using the *proseq2.0* pipeline (https://github.com/Danko-Lab/proseq2.0) to automate routine pre-processing, alignment, and coverage file generation. Paired-end FASTQ files were adapter-trimmed, quality filtered, and deduplicated based on UMI barcodes, then aligned to the reference genome using BWA. Aligned reads were converted to strand-specific coverage formats, reporting the 5′ end of nascent RNA reads.

