## Supplemental Information for "CpG island density predicts CBP/p300 dependency across 3D chromatin clusters"

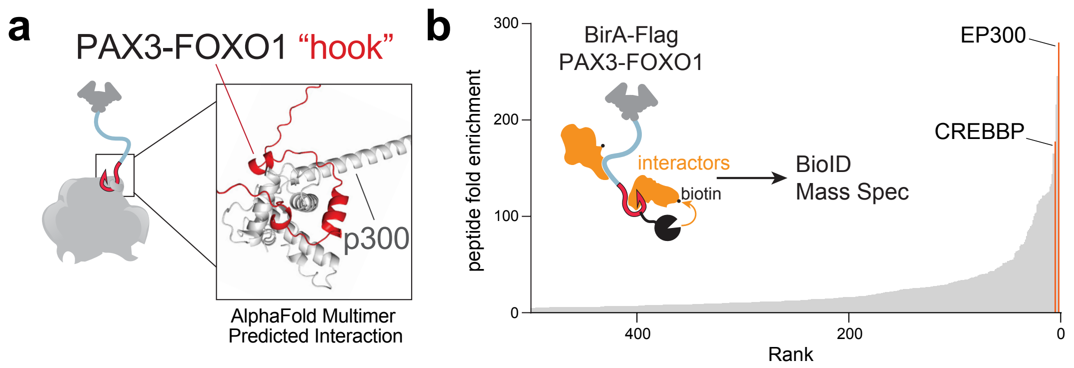


**Supplemental Figure 1. PAX3-FOXO1 recruits CBP/p300 to its activation domain, related to Figure 1**

**a.** Alphafold Multimer prediction of the interaction between the P3F activation domain “hook” and the KIX domain of p300. **b.** BioID mass spectrometry identification of CBP and p300 as top interactors with P3F, as previously described.

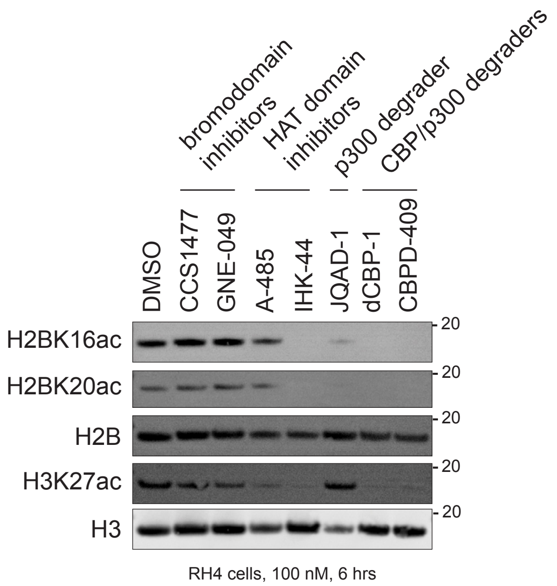


**Supplemental Figure 2. IHK-44 reduces histone acetylation comparable to CBP/p300 degraders, related to Figure 2**

Western blot of CBP/p300 bromodomain inhibitors (CCS1477, GNE-409), HAT domain inhibitors (A-485, IHK-44), selective p300 degrader (JQAD1), and dual CBP/p300 degraders (dCBP-1, CBPD-409) in RH4 cells after treatment at 100 nM for 6 h. H2B and H3 are shown as loading controls.

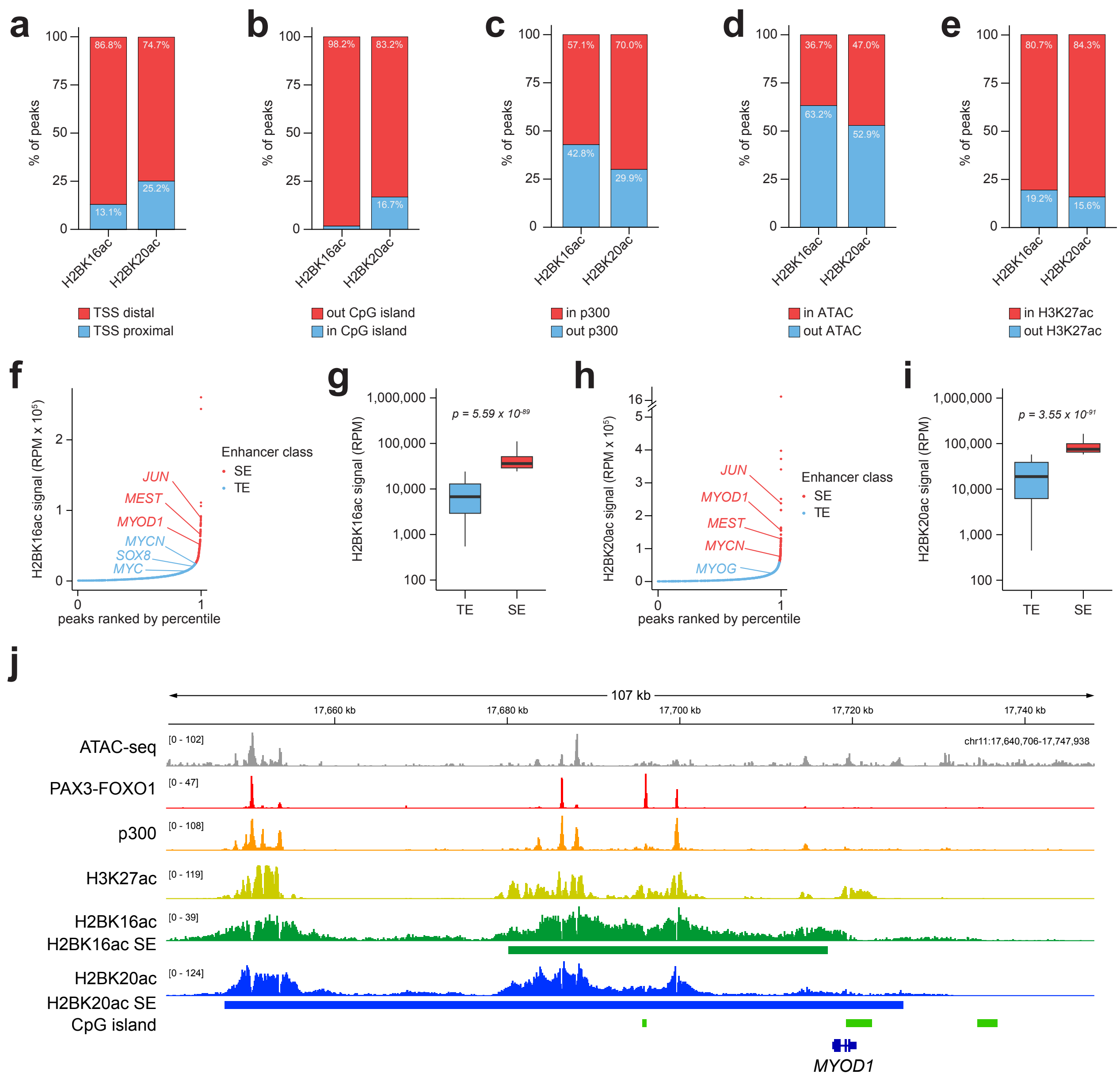


**Supplemental Figure 3. Baseline H2BNTKac landscape in FP-RMS, related to Figure 2**

**a-e.** Distribution of H2BK16ac and H2BK20ac ChIP-seq peaks in RH4 cells relative to transcription start sites (TSS) of gene promoters (**a**), CpG islands (**b**), p300 sites (**c**), ATAC-seq sites (**d**), and H3K27ac sites (**e**). **f.** Ranked ordered plot of H2BK16ac ChIP-seq signal (reads per million; RPM) of super-enhancers (SEs) and typical enhancers (TEs) over H2BK16ac sites in RH4 cells. **g.** Quantification of H2BK16ac ChIP-seq signal between TEs (blue) and SEs (red). Box plots represent median and quartiles, whiskers representing 1.5 x IQR. Statistical significance determined using a two-sided Wilcoxon rank-sum test. **h.** Ranked ordered plot of H2BK20ac ChIP-seq signal (RPM) of SEs and TEs over H2BK20ac sites in RH4 cells. **i.** Quantification of H2BK20ac ChIP-seq signal between TEs (blue) and SEs (red). Box plots represent median and quartiles, whiskers representing 1.5 x IQR. Statistical significance determined using a two-sided Wilcoxon rank-sum test. **j.** ChIP-seq signal at the *MYOD1* locus visualized in IGV.

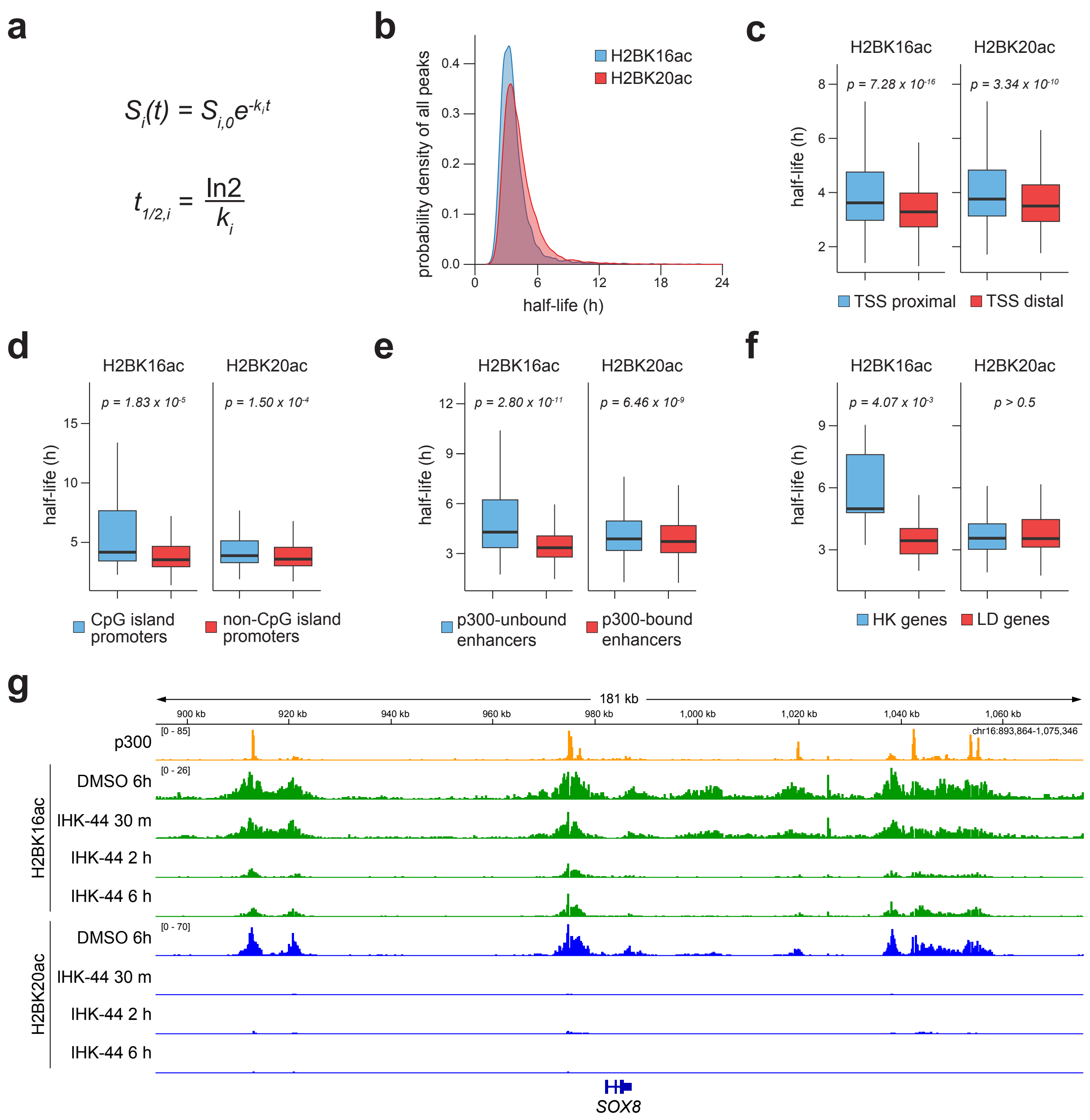


**Supplemental Figure 4. IHK-44 selectively reduces H2BNTKac at enhancers, related to Figure 2**

**a.** Formulas used for calculating histone acetylation decay rates and half-lives from integrated analysis of time-course treatment ChIP-seq datasets. **b.** Distribution of H2BK16ac and H2BK20ac ChIP-seq half-lives in RH4 cells. **c-f.** Distribution of H2BK16ac and H2BK20ac ChIP-seq half-lives in RH4 cells relative to transcription start sites (TSS) of gene promoters (**c**), CpG island-containing promoters (**d**), p300 sites (**e**), and gene classes (**f**). Box plots represent median and quartiles, whiskers representing 1.5 x IQR. Statistical significance determined using a two-sided Wilcoxon rank-sum test. **g.** ChIP-seq signal at the *SOX8* locus shows time-course reduction in H2BK16ac and H2BK20ac signal after IHK-44 treatment visualized in IGV.

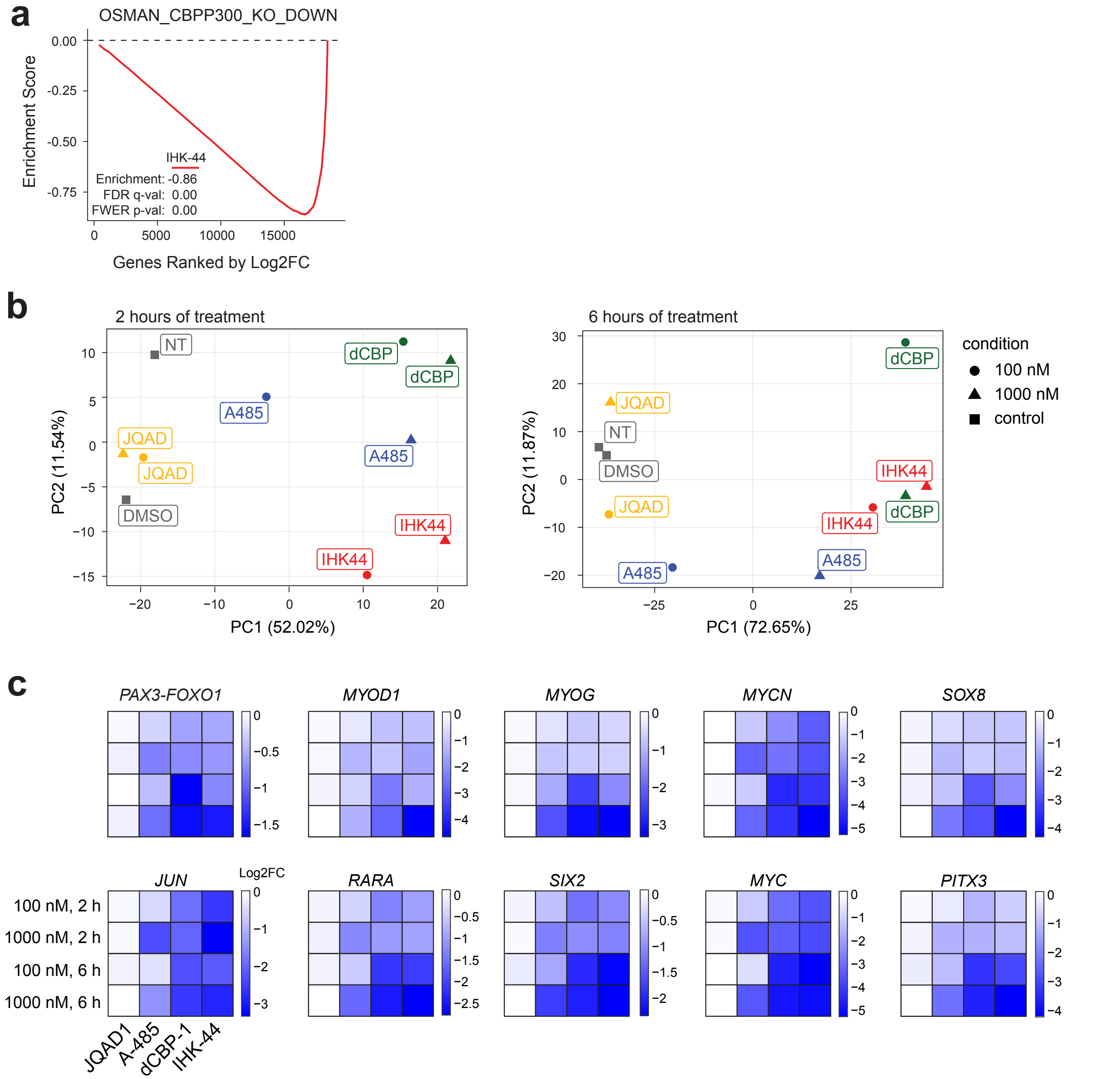


**Supplemental Figure 5. IHK-44 functionally mimics dual CBP/p300 degradation, related to Figure 3**

**a.** GSEA analysis of genes downregulated after CBP/p300 knockout in RH4 cells treated with 100 nM IHK-44 for 6 h. **b.** Principal component analysis of RNA-seq profiles following 2 h (left) and 6 h (right) treatment of RH4 cells at 100 nM or 1000 nM of JQAD1, A-485, dCBP-1, or IHK-44. **c.** Degree of downregulation in gene expression levels of FP-RMS core regulatory transcription factors (*PAX3-FOXO1, MYOD1, MYOG, MYCN, SOX8, JUN, RARA, SIX2, MYC, PITX3*) in RH4 cells treated with JQAD1, A-485, dCBP-1, or IHK-44.

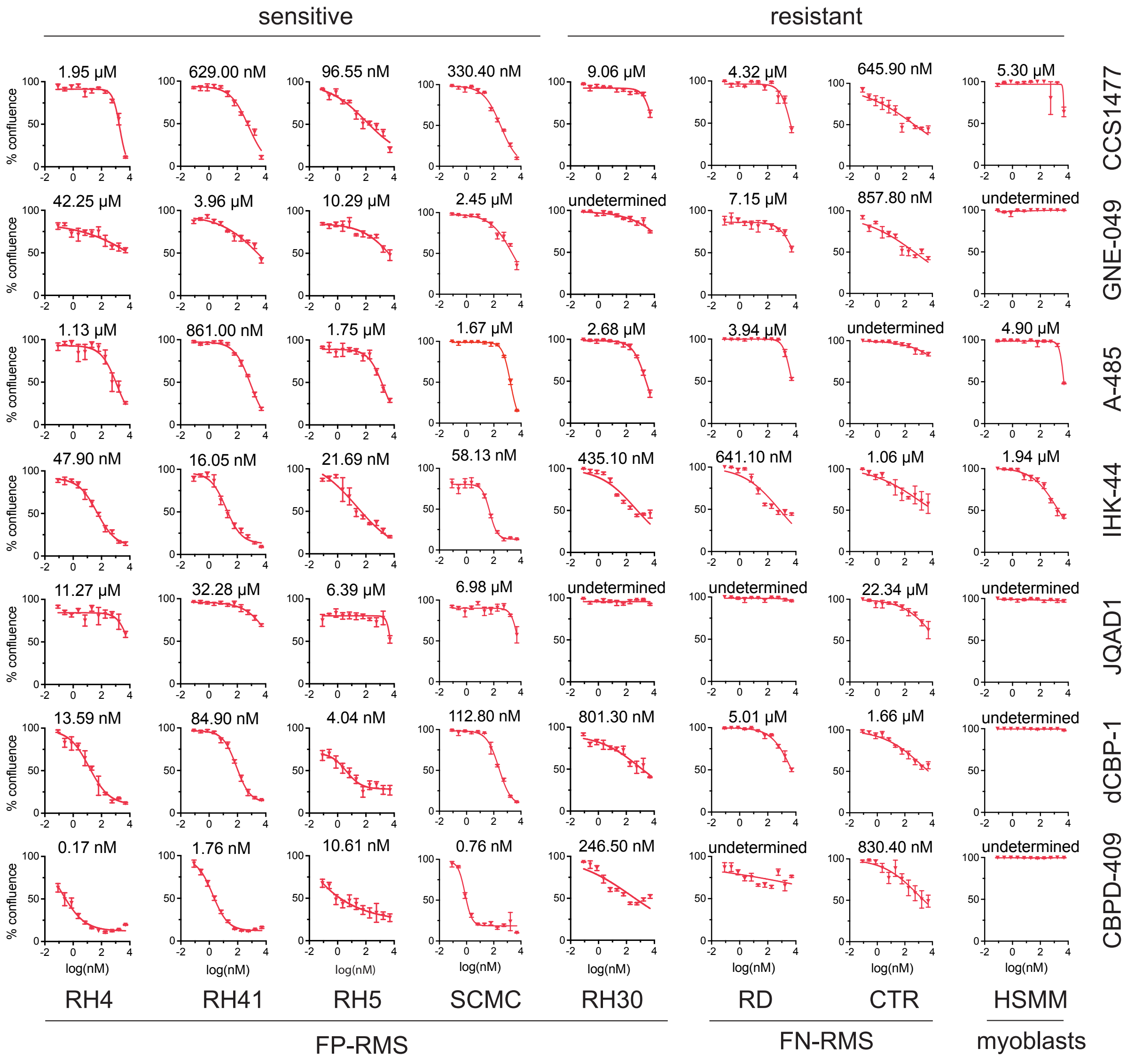


**Supplemental Figure 6. IHK-44 selectively reduces FP-RMS cell proliferation, related to Figure 5**

Dose-response curves of FP-RMS (RH4, RH41, RH5, SCMC, RH30), FN-RMS (RD, CTR), and normal myoblast (HSMM) cells treated with CBP/p300 bromodomain inhibitors (CCS1477, GNE-049), CBP/p300 HAT inhibitors (A-485, IHK-44), selective p300 degrader (JQAD1), or CBP/p300 degraders (dCBP-1, CBPD-409). $IC_{50}$ values are indicated for each treatment. Values represent mean ± SEM of *n = 3* technical replicates.

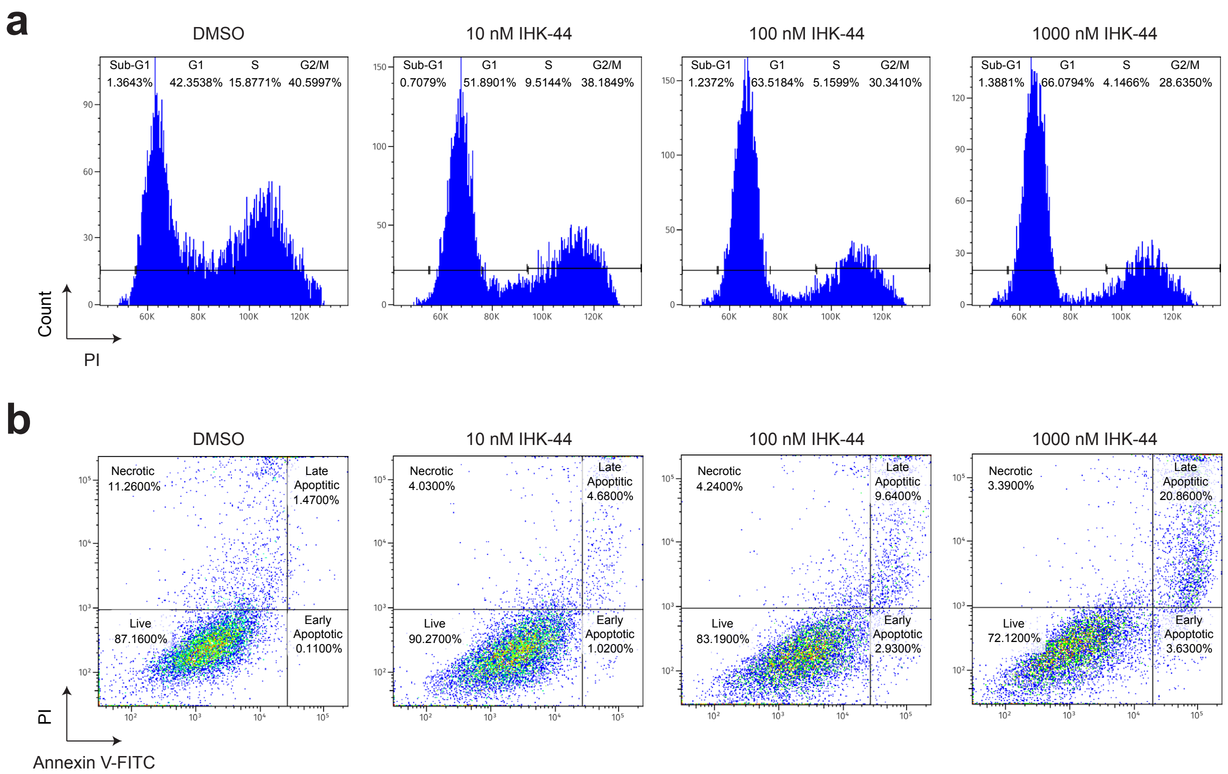


**Supplemental Figure 7. IHK-44 induces growth arrest in RH4 cells, related to Figure 5**
**a.** Cell-cycle arrest histograms of RH4 cells treated with increasing doses (10, 100, 1000 nM) of IHK-44 for 72 h. **b.** Apoptosis Annexin V/PI quadrants of RH4 cells treated with increasing doses of IHK-44 (10, 100, 1000 nM) for 72 h.

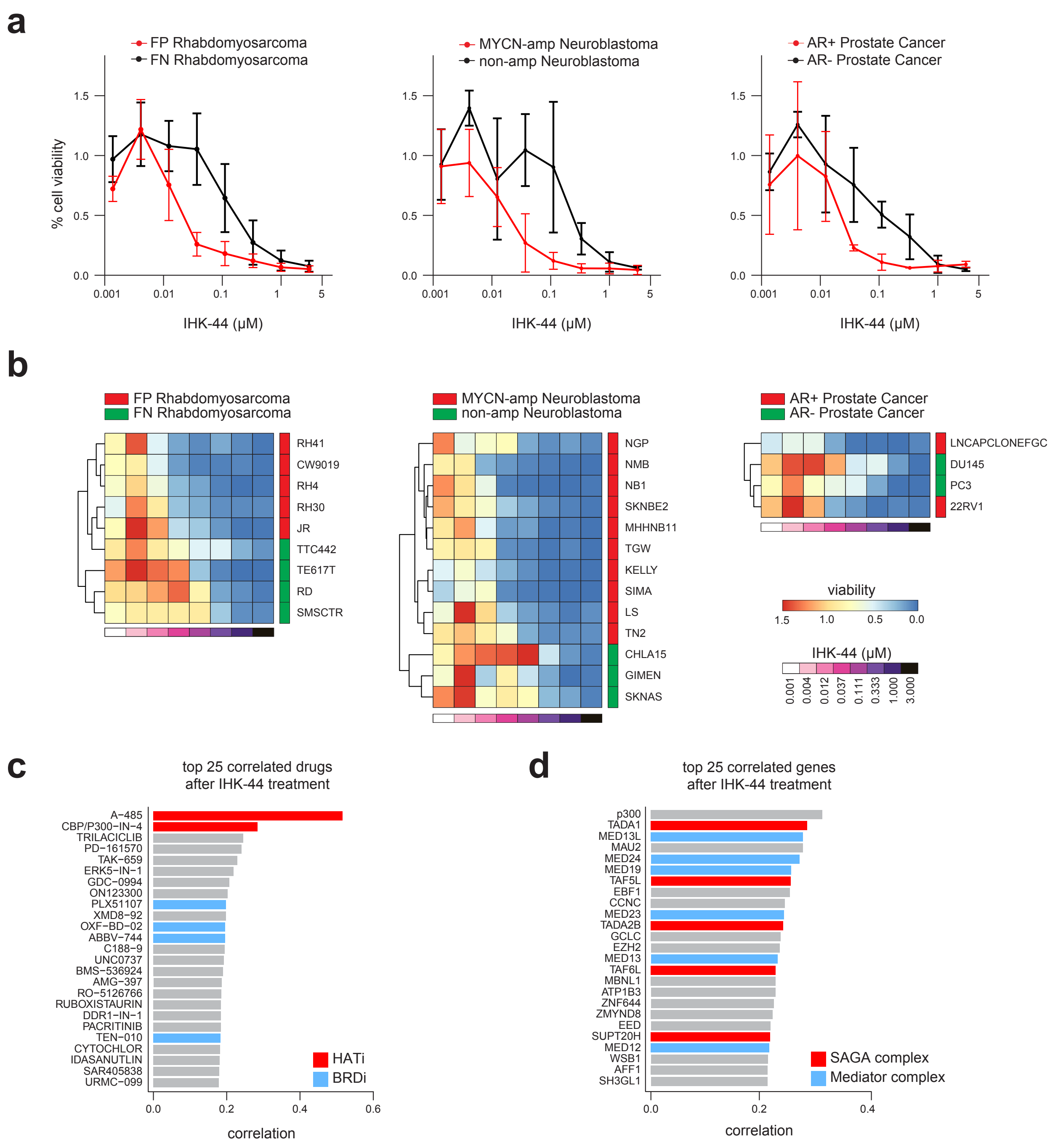


**Supplemental Figure 8. PRISM profiling reveals selective vulnerability of enhancer-addicted cancers to IHK-44, related to Figure 5**

**a.** Dose-response curves of cell viability of FP-RMS vs. FN-RMS, MYCN-amp NB vs. non-amp NB, and AR+ PCa vs. AR- PCa cell lines treated with IHK-44. Enhancer-addicted cancers (FP-RMS, MYCN-amp, AR+ PCa) are highlighted in red. Values represent mean ± SEM viability of all the cell lines in that subtype. **b.** Heatmaps depicting cell viability across cell lines from **a** within each lineage group. **c.** Top 25 drugs correlated with IHK-44 in the DepMap Repurposing dataset. Bromodomain inhibitors (BRDi) are highlighted in blue. HAT inhibitors (HATi) are highlighted in red. **d.** Top 25 genes correlated with IHK-44 in the DepMap CHRONOS dataset. Members of the SAGA complex (TADA1, TADA2B, TAF6L, SUPT20H) are highlighted in red. Members of the Mediator complex (MED13L, MED24, MED19, MED23, MED13, MED12) are highlighted in blue.

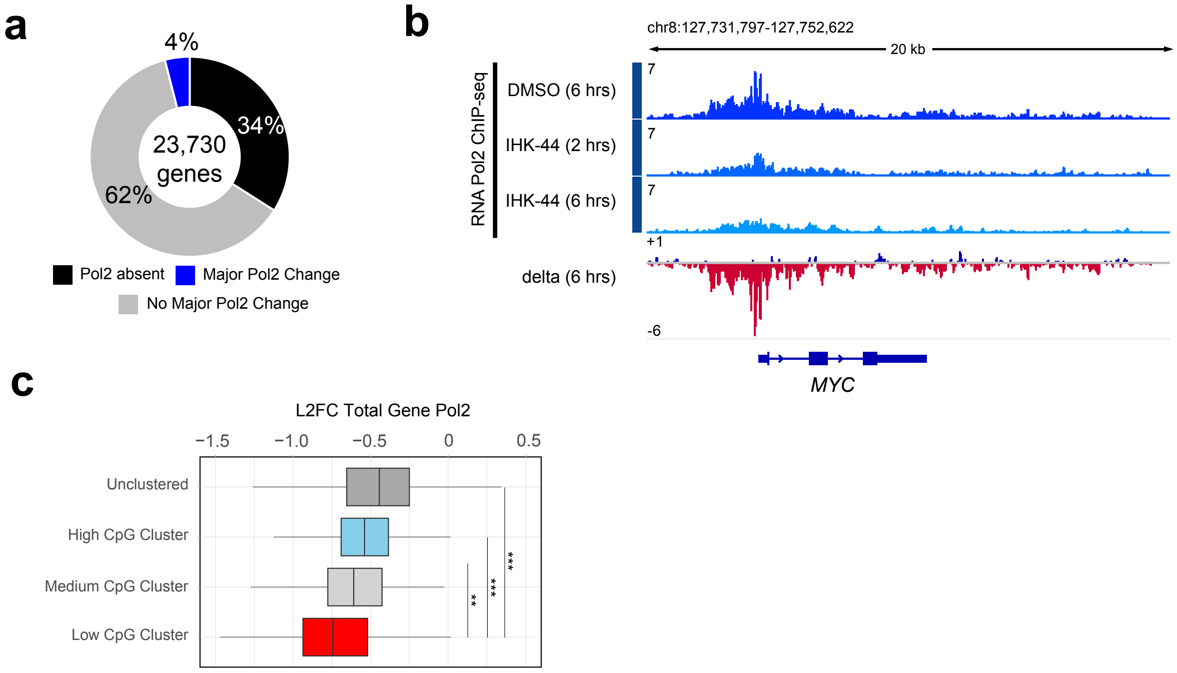


**Supplemental Figure 9. IHK-44 treatment induces substantial RNA Pol2 loss at a select subset of critical genes, related to Figure 6**

**a.** Donut plot showing the percentages of genes with trivial Pol2, no major change and major Pol2 disruption following IHK-44 treatment for 6 hours. **b.** RNA Pol2 ChIP-seq at the MYC locus across DMSO, IHK-44 (2h) and IHK-44 (6h) timepoints visualized with IGV. **c.** Boxplots of L2FC of total RNA Pol2 following IHK-44 treatment at 6 hours, split between genes not in Pol2 HiChIP clusters and those genes in high, medium and low frequency CpG island clusters (box plots of median and quartiles, whiskers showing 1.5 × inter-quartile ranges; Mann-Whitney U test for significance)

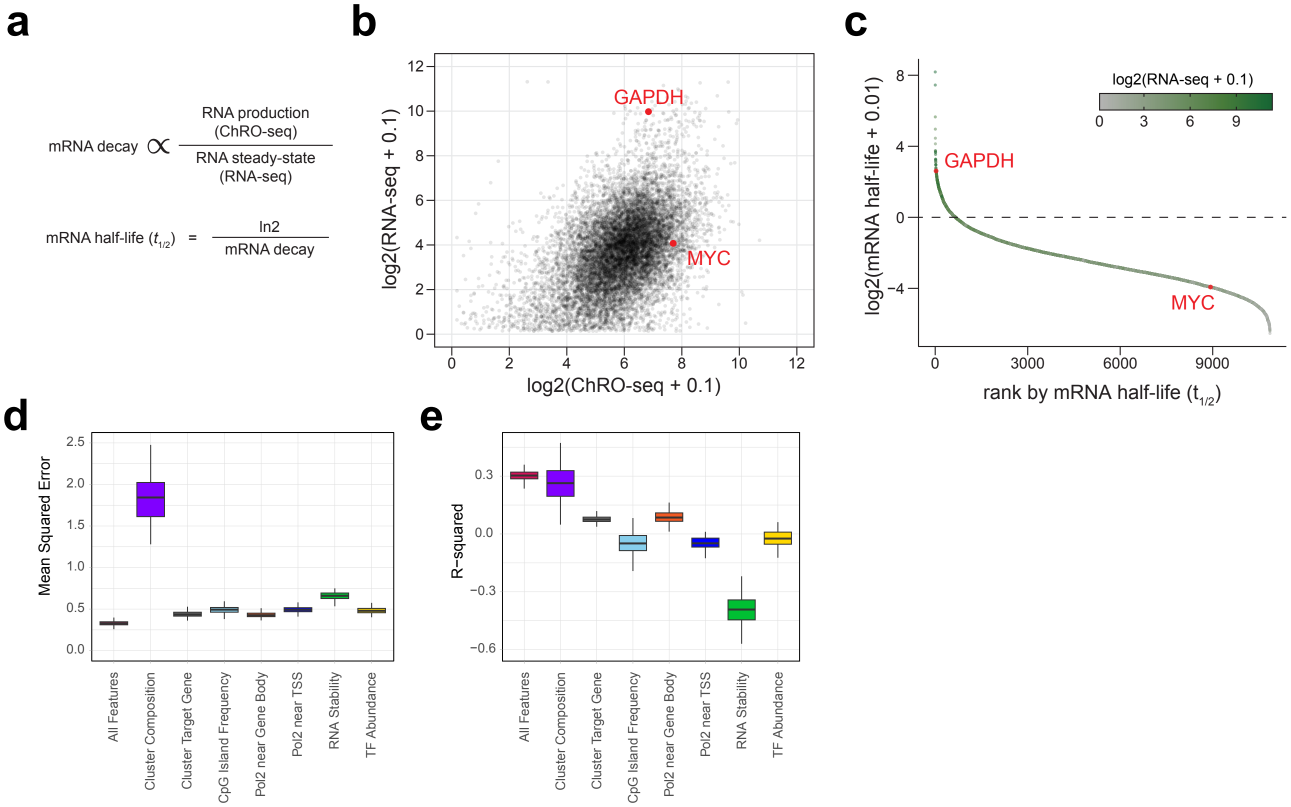


**Supplemental Figure 10. RNA stability separates classes of genes and provides predictive power in concert with other epigenomic features, related to Figure 7**

**a.** Formulas for calculating RNA half-life from ChRO-seq and RNA-seq TPM values. **b.** Scatter plot of genes by log2 ChRO-seq and log2 RNA-seq, with housekeeping gene GAPDH and transcription factor MYC highlighted in red. **c.** Rank plot of genes by mRNA half-life, with GAPDH and MYC highlighted in red. **d.** Mean squared error across 100 runs of random forest regression models, split by the features included in the training (box plots of medians and quartiles, whiskers showing 1.5 × inter-quartile ranges) **e.** $R^2$ values from across 100 runs of random forest regression models, split by the features included in the training (box plots of medians and quartiles, whiskers showing 1.5 × inter-quartile ranges)

**General Chemistry**

The chemicals were purchased from various vendors (Sigma-Aldrich, Fisher, Oakwood, Combi Blocks, and AA Blocks) and used as received, unless specified. All syntheses were conducted with anhydrous solvents under an atmosphere of argon or nitrogen, using oven-dried glassware and employing standard techniques for handling air-sensitive materials unless otherwise noted. All solvents were anhydrous and stored under an argon or nitrogen atmosphere before use.

Analytical thin-layer chromatography (TLC) was performed on Kieselgel 60 F254 glass plates, precoated with a 0.25 mm thickness of silica gel. TLC plates were visualized using UV light and staining reagents, including iodine, anisaldehyde, and permanganate (KMnO4). Hydrogenation reactions were performed using an atmospheric balloon. Purification of all intermediate and final products was achieved using the CombiFlash Nextgen 100 (Teledyne Isco). Normal-phase chromatography was performed on a Teledyne ISCO CombiFlash NextGen300 system using Teledyne RediSep normal-phase silica cartridges with an average particle size of 35–70 µm, while RediSep Gold® C18 Reversed Phase Columns (20-40 microns) were used for reverse-phase chromatography. The $^{1}$H, $^{13}$C NMR nuclear magnetic resonance (NMR) spectra were recorded at 25 °C on 400 MHz on a Bruker Avance Neo NanoBay 400 spectrometer with standard pulse sequences or Bruker AVANCE NMR spectrometer operating at 500 MHz. Chemical shifts were reported in δ units, part per million, with reference to the residual solvent peak $CDCl_3$ (δ 7.26) and DMSO-$d_6$ (δ 2.50) for $^{1}$H and $CDCl_3$ (δ 77.3) and DMSO-$d_6$ (δ 39.5) for $^{13}$C NMR spectra. Coupling constants are reported in hertz (Hz). The following abbreviations (or a combination thereof) are used to describe splitting patterns: s, singlet; d, doublet; t, triplet; q, quartet; pent, pentet; m, multiplet; br, broad. All compounds were of 95% purity or higher, unless otherwise noted, as measured by analytical reverse-phase HPLC. High-resolution mass spectra were recorded on an Agilent 1290 Infinity II Series 6230B TOF LC/MS. Detection methods included diode-array (DAD) at 210 and 254 nm and positive/negative electrospray ionization (ESI), with a mass range of 25–20 000 *m*/*z*. The MS detector was configured to a mass range of 100 to 1700 m/z , with nitrogen used as the nebulizer gas. High-resolution acquisition-rate mass spectra (MS mode) were acquired in electrospray mode by scanning at a rate up to 40 spectra/s. The system can maintain a 2 ppm mass accuracy/stability within 2 °C drift per hour. Data acquisition was performed with MassHunter Walkup software. Unless stated, all methods use an Agilent InfinityLab Poroshell 120 EC-C18 column, dimensions 4.6 × 50 mm, 2.7 µm, fitted with Poroshell 120 EC-C18, 2.1 mm, 1.9 µm guard. Mobile phase A was 0.1% TFA in $H_2O$; mobile phase B was 0.1% TFA in $CH_3CN$.

For LC-MS, the following methods and conditions were used.
Method A: Reverse-phase HPLC was carried out with a flow rate of 0.4 mL/min, at 55 °C, using positive ESI mode. Injection volume = 1 µL. The gradient conditions used were 5% mobile phase B for 0.2 min, then a gradient from 5% to 95% over 2.0 min, and then held at 95% mobile phase B for 0.45 min.
Method B: Reverse-phase HPLC was carried out with a flow rate of 0.4 mL/min, at 55 °C, using positive ESI mode. Injection volume = 1 µL. The gradient conditions used were 40% mobile phase B for 0.2 min, then a gradient of 40–95% mobile phase B over 2.5 min, and then held at 95% mobile phase B for 0.5 min.

### General Procedures

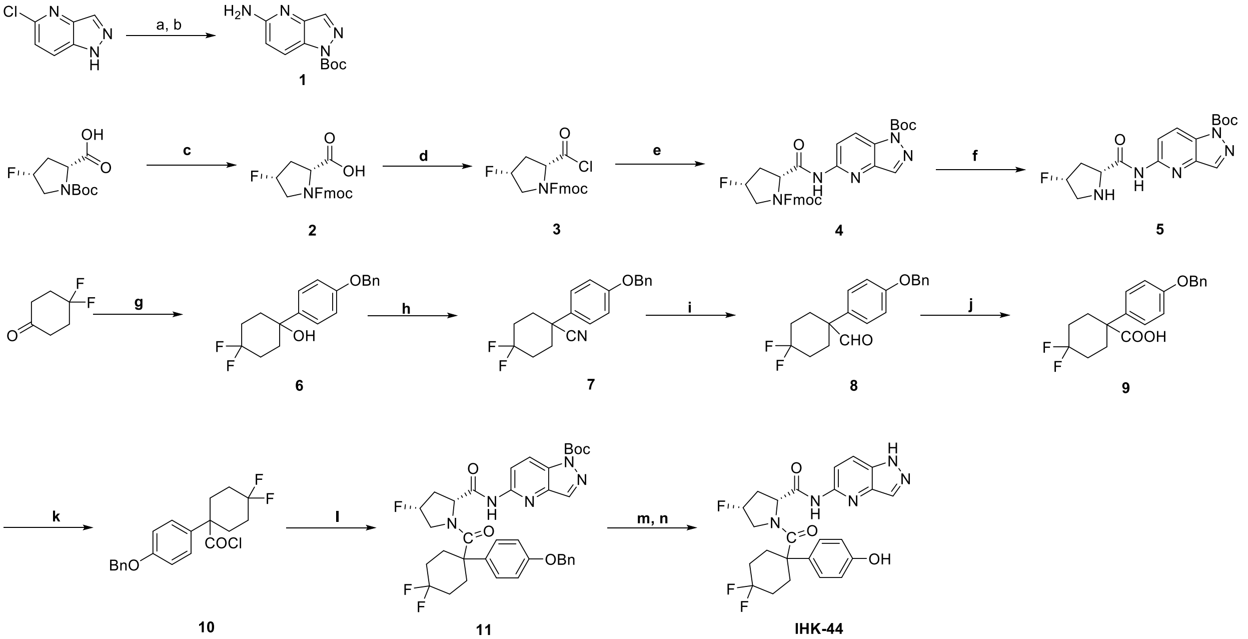

**Reagents and conditions**: (a) LHMDS, $Pd_2(dba)_3$, XPhos, THF, reflux; (b) $Boc_2O$, TBAF, 0 °C, 72% (two steps); (c) (i) 4 N HCl/1,4-dioxane, rt; (ii) $NaHCO_3$, FmocCl, 1,4-dioxane/$H_2O$, rt, 85%; (d) $SOCl_2$, DMF, DCM, rt to 40 °C; (e) **1**, DIPEA, DCM, 0 °C to rt, 68%; (f) piperidine, DMF, 0 °C to rt, 80%; (g) 4-Benzyloxybromobenzene, nBuLi (2.5 M in Hexane), THF, -78 °C to rt; (h) $InBr_3$, TMSCN, DCM, rt, 85%; (i) DIBALH, toluene, rt; (j) $NaClO_2$, 2-methyl-2-butene, $KH_2PO_4$, tBuOH/$H_2O$, rt, 50% (two steps) (k) $SOCl_2$, DMF, DCM, rt to 40 °C; (l) **5**, DIPEA, DMF, DCM, 0 °C to rt; 41% (two steps) (m) $H_2$, Pd in C (10%), EtOH:THF (1:1) rt; (n) TFA, DCM, rt, 94%

#### ***tert*-Butyl 5-amino-1*H*-pyrazolo[4,3-b]pyridine-1-carboxylate (1)**

To a solution of commercially available 5-chloro-1H-pyrazolo[4,3-b]pyridine (5 g, 32.8 mmol) dissolved in THF (34 mL) was added $Pd_2(dba)_3$ (772 mg, 0.84 mmol), XPhos (622 g, 1.3 mmol), LHMDS (1.09 mol/L solution in THF, 68 mL, 74 mmol), and stirred under reflux for 6 h. The reaction mixture was allowed to cool at room temperature and then stirred overnight. The reaction mixture was cooled to 0°C, and $BOC_2O$ (7.62 g, 34.9 mmol) was added in small portions. The mixture was stirred for 40 min at the same temperature. Then, TBAF (95 mmol) was added, and the resultant mixture was stirred at 0°C for 1h. Water was added to the mixture at 0°C to quench the reaction. The reaction mixture was extracted with ethyl acetate three times, the organic layers were combined, washed with brine, and dried over anhydrous sodium sulfate. The solvent was evaporated under reduced pressure to obtain a crude extract. The crude extract was purified by silica gel chromatography (hexane/ethyl acetate) to obtain **1** (5.51 g, 72%) as a pale yellow solid. $^1H$ NMR (400 MHz, $CDCl_3$) δ 8.00 (d, $J$ = 9.1 Hz, 1H), 7.90 (s, 1H), 6.61 (d, $J$ = 9.1 Hz, 1H), 5.12 (s, 2H), 1.56 (s, 9H).

#### **(2*R*,4*R*)-1-(((9*H*-fluoren-9-yl)methoxy)carbonyl)-4-fluoropyrrolidine-2-carboxylic acid (2)**

To a solution of (2*R*,4*R*)-1-(*tert*-butoxycarbonyl)-4-fluoropyrrolidine-2-carboxylic acid (1.17 g, 5 mmol) was added 4M HCl in dioxane (11.65 mL) and stirred for 4h at room temperature. The reaction mixture was concentrated under reduced pressure to obtain a solid, which was dissolved in water (23 mL), ice cooled, and supplemented with $NaHCO_3$ (2.09 g, 24.9 mmol), 1,4-dioxane (23.25 mL), and 9-fluorenylmethylchloroformate (4.00 g, 15.4 mmol), and stirred at room temperature overnight. Water was added to the reaction mixture, which was then washed with diethyl ether twice, and the aqueous layer was acidified with 1N HCl. The reaction mixture was extracted with chloroform three times, dried over anhydrous sodium sulfate, filtered, and concentrated under reduced pressure. The obtained residue was purified by silica gel column chromatography (dichloromethane/10% methanol in dichloromethane) to obtain **2** (1.51 g, 85%) as a colorless solid. $^1$H NMR (400 MHz, DMSO) δ 12.77 (s, 1H), 7.89 (t, *J* = 7.0 Hz, 2H), 7.66 (dd, *J* = 14.7, 7.9 Hz, 2H), 7.42 (dt, *J* = 9.6, 4.6 Hz, 2H), 7.33 (q, *J* = 8.6 Hz, 2H), 5.51 – 5.04 (m, 1H), 4.60 – 4.06 (m, 4H), 3.88 – 3.39 (m, 2H), 2.77 – 2.06 (m, 2H).

***tert*-Butyl 5-((2*R*,4*R*)-1-(((9*H*-fluoren-9-yl)methoxy)carbonyl)-4-fluoropyrrolidine-2-carboxamido)-1*H*-pyrazolo[4,3-b]pyridine-1-carboxylate (4)**

To a solution of **2** (1.11 g, 3.1 mmol) dissolved in dichloromethane (18 mL), thionyl chloride (2.27 mL) and DMF (31 μL) were added, and stirred at room temperature for 3h and then 40 ºC for 30 minutes. The reaction solution was concentrated under reduced pressure, dissolved in 3 mL dichloromethane, and supplemented with hexane (30 mL). The precipitated solid was collected by filtration and dried to obtain acid chloride **3** as a pale yellow solid.

To a mixture of **1** (731 mg, 3.12 mmol), DIPEA (706.5 μL, 4.1 mmol), and dichloromethane (9 mL) was added crude acid chloride **3** dissolved in dichloromethane (9 mL) under ice cooling, followed by stirring at room temperature for 3h. Then, 1mol/L HCl was added to the reaction solution, and the reaction mixture was extracted with dichloromethane three times, dried over anhydrous sodium sulfate, filtered, and concentrated under reduced pressure. The obtained residue was purified by silica gel column chromatography (hexane/ethyl acetate) to obtain **4** (1.18 g, 68% yield) as a colorless solid. $^1$H NMR (400 MHz, DMSO) δ 11.04 – 10.60 (m, 1H), 8.58 – 8.24 (m, 3H), 7.91 (d, *J* = 7.4 Hz, 1H), 7.78 (s, 1H), 7.70 (s, 1H), 7.58 (d, *J* = 7.5 Hz, 1H), 7.50 – 7.19 (m, 3H), 7.05 (d, *J* = 8.0 Hz, 1H), 5.50 – 5.16 (m, 1H), 4.81 – 4.56 (m, 1H), 4.41 – 4.08 (m, 3H), 3.90 – 3.64 (m, 2H), 2.78 – 2.54 (m, 1H), 2.45 – 2.22 (m, 1H), 1.66 (s, 9H).

***tert*-Butyl 5-((2*R*,4*R*)-4-fluoropyrrolidine-2-carboxamido)-1*H*-pyrazolo[4,3-b]pyridine-1-carboxylate (5)**

To a solution of **4** (150 mg, 0.26 mmol) dissolved in dimethylformamide (5.2 mL), piperidine (257 μL, 2.6 mmol) was added under ice-cooling, followed by stirring under ice-cooling for 15 minutes and at room temperature for 15 minutes. Water was added to the reaction solution, and the reaction mixture was extracted with ethyl acetate three times. The organic layer was washed with saturated brine and dried over anhydrous sodium sulfate. Then, the aqueous layers were combined, extracted with dichloromethane twice, and the organic layer was dried over anhydrous sodium sulfate. Both organic layers were combined, filtered, and concentrated under reduced pressure. The obtained residue was purified by silica gel column chromatography (ethyl acetate/methanol) to obtain **5** (72.5 mg ,80%) as a colorless solid. $^1$H NMR (400 MHz, $CDCl_3$) δ 10.61 (s, 1H), 8.56 – 8.10 (m, 3H), 5.53 – 5.06 (m, 1H), 4.97 – 4.53 (m, 1H), 4.29 – 3.81 (m, 1H), 3.81 – 3.40 (m, 1H), 3.04 – 2.80 (m, 1H), 2.79 – 2.42 (m, 2H), 1.71 (s, 9H).

**1-(4-(benzyloxy)phenyl)-4,4-difluorocyclohexan-1-ol (6)**

To a solution of 1-(benzyloxy)-4-bromobenzene (1.58 g, 6 mmol) in THF (12 mL) was added n-butyllithium (2.5 M in hexane, 3.12 mL, 7.8 mmol) at -78 ºC and stirred for 30 minutes. 4,4-difluorocyclohexan-1-one (1.13 g, 8.4 mmol) was dissolved in THF (8.4 mL) and added slowly to the reaction mixture at the same temperature, and the mixture was stirred for 2 h. The reaction mixture is then heated to room temperature, and the reaction progress is monitored by TLC and continued until the starting material is fully consumed. The reaction mixture was quenched with water and acidified with 1 M HCl. The reaction mixture was acidified with 1 M HCl and extracted with ethyl acetate three times; the organic layers were combined, dried over anhydrous sodium sulfate, filtered, and concentrated under reduced pressure. The obtained residue was purified by silica gel column chromatography (hexane/ethyl acetate) to obtain **6** (955 mg, 50% yield) as a colorless solid.$^{1}$H NMR (400 MHz, DMSO) δ 7.51 – 7.23 (m, 7H), 6.96 (dq, *J* = 8.8, 2.4 Hz, 2H), 5.08 (d, *J* = 5.1 Hz, 3H), 2.33 – 2.07 (m, 2H), 1.91 (q, *J* = 9.8 Hz, 4H), 1.73 (d, *J* = 13.4 Hz, 2H).

**1-(4-(benzyloxy)phenyl)-4,4-difluorocyclohexane-1-carbonitrile (7)**

Compound **6** (1.6 g, 5.03 mmol) was dissolved in DCM (10 mL). To this solution, indium bromide (178 mg, 10 mmol%) and trimethylsilyl cyanide (1.26 mL, 10.1 mmol) dissolved in DCM (10 mL) were added dropwise, and the mixture was stirred at room temperature for 30 minutes. The reaction mixture was concentrated, and the crude extract was purified by silica gel column chromatography (hexane/ethyl acetate) to obtain **7** (1.4 g, 85%) as a colorless oil. $^{1}$H NMR (400 MHz, $CDCl_3$) δ 7.49 – 7.31 (m, 7H), 7.02 (d, *J* = 8.3 Hz, 2H), 5.09 (s, 2H), 2.43 – 2.02 (m, 8H).

**1-(4-(benzyloxy)phenyl)-4,4-difluorocyclohexane-1-carboxylic acid (9)**

To a solution of **7** (1.2 g, 3.67 mmol) in toluene (11 mL), diisobutylaluminum hydride (1.2 M in toluene, 4.58 mL, 5.5 mmol) was added, and the mixture was stirred for 20 minutes. After that, a saturated aqueous solution of l-(+)-potassium sodium tartrate was added to quench the reaction mixture, which was stirred at room temperature for 5 min. The mixture was extracted with ethyl acetate three times, and the organic layers were combined, washed with 1 mol/L HCl, and with saturated brine. The organic layers were dried over anhydrous sodium sulfate and concentrated to obtain crude 1-(4-(benzyloxy)phenyl)-4,4-difluorocyclohexane-1-carbaldehyde **8** as a colorless oil. The crude mixture was then dissolved in tert-butyl alcohol (18 mL) and water (3.67 mL). Under ice cooling, to the reaction mixture, 2-methyl-2-butene (1.94 mL, 18.35 mmol), sodium dihydrogenphosphate (880 mg, 7.24 mmol), and sodium chlorite (664 mg, 7.24 mmol) were sequentially added and stirred at room temperature for 3h. Then, 1 mol/L HCl was added to quench the reaction, the reaction mixture was extracted with ethyl acetate three times, the organic layers were combined, washed with saturated brine, and dried over sodium sulfate. The crude extract was purified by silica gel column chromatography (hexane/ethyl acetate) to obtain **9** (635.6 mg, 50%) as a white solid.$^{1}$H NMR (400 MHz, DMSO) δ 12.66 (s, 1H), 7.57 – 7.27 (m, 7H), 7.11 – 6.83 (m, 2H), 5.09 (s, 2H), 2.47 – 2.32 (m, 2H), 2.14 – 1.73 (m, 6H).

***tert*-Butyl 5-((2*R*,4*R*)-1-(1-(4-(benzyloxy)phenyl)-4,4-difluorocyclohexane-1-carbonyl)-4-fluoropyrrolidine-2-carboxamido)-1H-indazole-1-carboxylate 11**

To a solution of 9 (793 mg, 2.29 mmol) in DCM (14 mL) was added thionyl chloride (1.66 mL, 22.9 mmol). A catalytic amount of DMF (25 µL) was added to the reaction mixture dropwise. The reaction mixture was stirred for 3 hours at room temperature, then heated to 40 ºC and stirred for an additional 30 minutes. The reaction solution was concentrated under reduced pressure, dissolved in 3 mL dichloromethane, and supplemented with hexane (30 mL). The precipitated solid was collected by filtration and dried to obtain acid chloride **10** (534 mg) as a pale yellow solid. To the solution of **5** (268 mg, 0.77 mmol), DIPEA (174 µL) in DCM (2 mL) was cooled down to 0 °C, followed by the dropwise addition of a solution of acid chloride **10** (280 mg, 0.77 mmol) dissolved in DCM (2 mL). The reaction mixture was stirred at 0 °C, then heated to room temperature, and stirred until the starting materials were fully consumed. The reaction mixture was quenched with water, then with 1 M HCl. The reaction mixture was extracted with dichloromethane three times, combined with the organic layers, washed with brine, and dried over sodium sulfate. The crude mixture was evaporated and purified by silica gel column chromatography (hexane/ethyl acetate) to obtain **11** (635 mg, 41%) as a yellowish solid. 1H

NMR (400 MHz, DMSO) δ 10.80 (s, 1H), 8.57 – 8.25 (m, 3H), 7.57 – 7.22 (m, 7H), 7.04 (d, J = 8.0 Hz, 2H), 5.11 (d, J = 11.6 Hz, 2H), 5.05 – 4.74 (m, 2H), 3.16 – 2.92 (m, 1H), 2.47 – 1.81 (m, 11H), 1.66 (s, 9H). [19]F NMR (376 MHz, DMSO) δ -89.80, -90.42, -98.00, -98.62, -173.39.

**(2*R*,4*R*)-1-(4,4-difluoro-1-(4-hydroxyphenyl)cyclohexane-1-carbonyl)-4-fluoro-N-(1*H*-indazol-5-yl)pyrrolidine-2-carboxamide, IHK-44**

To a solution of IHK-45 (118.6 mg, 0.18 mmol) dissolved in ethanol (3.5 mL) and THF (3.5 mL), 10% Pd in C (18.62 mg, 10 mol%) was added. The reaction was stirred under a hydrogen atmosphere ($H_2$ balloon) at room temperature for 18 h. The crude reaction mixture was filtered through a pad of Celite, and the filtrate was concentrated under reduced pressure to afford the crude product (91 mg). The crude product was dissolved in DCM (2 mL), and trifluoroacetic acid (2 mL) was added. The reaction mixture was stirred at room temperature for 1 hour. The crude reaction mixture was evaporated and purified by silica gel column chromatography (hexane/ethyl acetate) to obtain **IHK-44** (75 mg, 94%) as a colorless solid. LCMS: LC/MS (ESI) m/z: 488.36 $(M+H)^+$. [1]H NMR (400 MHz, DMSO) δ 13.27 (s, 1H), 10.46 (s, 1H), 9.46 (s, 1H), 8.23 – 7.93 (m, 3H), 7.16 (d, *J* = 8.1 Hz, 2H), 6.79 (d, *J* = 8.0 Hz, 2H), 5.06 (d, *J* = 53.7 Hz, 1H), 4.79 (d, *J* = 9.6 Hz, 1H), 3.19 – 2.92 (m, 1H), 2.48 – 1.85 (m, 10H), 1.79 – 1.57 (m, 1H). [19]F NMR (376 MHz, DMSO) δ -89.79, -90.40, -97.96, -98.58, -173.57.

[1]H NMR of compound IHK-44

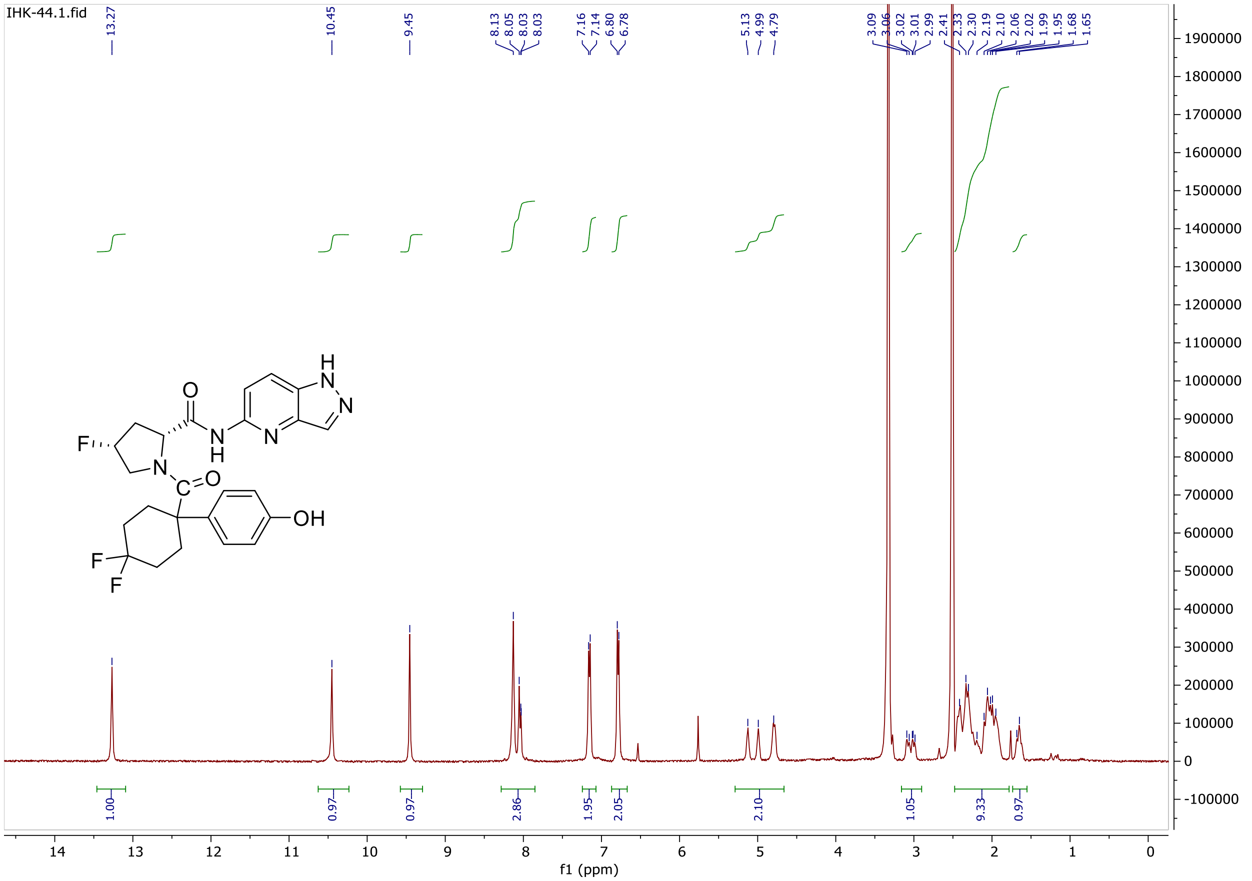
